# Multiscale analysis of single and double maternal-zygotic Myh9 and Myh10 mutants during mouse preimplantation development

**DOI:** 10.1101/2020.09.10.291997

**Authors:** Markus Frederik Schliffka, Anna-Francesca Tortorelli, Özge Özgüç, Ludmilla de Plater, Oliver Polzer, Diane Pelzer, Jean-Léon Maître

## Abstract

During the first days of mammalian development, the embryo forms the blastocyst, the structure responsible for implanting the mammalian embryo. Consisting of an epithelium enveloping the pluripotent inner cell mass and a fluid-filled lumen, the blastocyst results from a series of cleavages divisions, morphogenetic movements and lineage specification. Recent studies identified the essential role of actomyosin contractility in driving the morphogenesis, fate specification and cytokinesis leading to the formation of the blastocyst. However, the preimplantation development of contractility mutants has not been characterized. Here, we generated single and double maternal-zygotic mutants of non-muscle myosin-II heavy chains (NMHC) to characterize them using multiscale imaging. We find that Myh9 (NMHC II-A) is the major NMHC during preimplantation development as its maternal-zygotic loss causes failed cytokinesis, increased duration of the cell cycle, weaker embryo compaction and reduced differentiation, whereas Myh10 (NMHC II-B) maternal-zygotic loss is much less severe. Double maternal-zygotic mutants for Myh9 and Myh10 show a much stronger phenotype, failing most attempts of cytokinesis. We find that morphogenesis and fate specification are affected but nevertheless carry on in a timely fashion, regardless of the impact of the mutations on cell number. Strikingly, even when all cell divisions fail, the resulting single-celled embryo can initiate trophectoderm differentiation and lumen formation by accumulating fluid in increasingly large vacuoles. Therefore, contractility mutants reveal that fluid accumulation is a cell-autonomous process and that the preimplantation program carries on independently of successful cell division.

## Introduction

During embryonic development, cells execute their genetic program to build organisms with the correct cell fate, shape, position and number. In this process, the coordination between cell proliferation, differentiation and morphogenesis is crucial. In the mouse, the early blastocyst consists of 32 cells distributed among the trophectoderm (TE) and inner cell mass (ICM) with squamous TE cells enveloping the ICM and a lumen called the blastocoel [1,2]. Starting from the zygote, the early blastocyst forms after 5 cleavage divisions, 3 morphogenetic movements and 2 lineage commitments [3–5]. Differentiation and morphogenesis are coupled by the formation of an apical domain after the 3^rd^ cleavage, which acts as a signaling platform guiding cell division orientation [6,7], contact rearrangement [8] and differentiation [9]. Morphogenesis and cleavages are concomitant and appear synchronized with compaction starting after the 3^rd^ cleavage, internalization after the 4^th^ and lumen opening after the 5^th^ [3]. Beyond this apparent coordination, the existence of a coupling between cleavages and morphogenesis requires further investigations.

Actomyosin contractility is a conserved instrument driving animal morphogenesis [10,11] and cytokinesis [12]. Recent studies have suggested key contributions of actomyosin contractility during all the morphogenetic steps leading to the formation of the blastocyst [3,5]. During the 8-cell stage, increased contractility at the cell-medium interface pulls blastomeres together and compacts the embryo [13]. Also, cells form a domain of apical materials that inhibits actomyosin contractility [8,14]. During the 4^th^ cleavage, the asymmetric division of this domain leads sister cells to exhibit distinct contractility [8,15]. This causes the most contractile blastomeres to internalize and adopt ICM fate, while weakly contractile cells are stretched at the surface of the embryo and become TE [8,16]. When blastocoel fluid starts to accumulate, these differences in contractility between ICM and TE cells guide the fluid away from ICM-ICM cell-cell contacts [17]. Finally, contractility has been proposed to control the size of the blastocoel negatively by increasing the tension of the TE and positively by reinforcing cell-cell adhesion via mechanosensing [18]. Together, these findings highlight actomyosin as a major engine powering blastocyst morphogenesis [19].

To change the shape of animal cells, myosin motor proteins contract a network of cross-linked actin filaments, which can be tethered to the plasma membrane, adherens junctions and/or focal adhesions [20]. This generates tension, which cells use to change shape and tissue topology [21,22]. Among myosin motors, non-muscle myosin II are key drivers of cell shape changes [23]. Non-muscle myosin II complexes assemble from myosin regulatory light chains, myosin essential light chains and non-muscle myosin heavy chains (NMHC) [24]. NMHC are responsible for generating the power stroke and exist in 3 distinct paralogs in the mouse and human: NMHC II-A, II-B and II-C, encoded by the genes Myh9/MYH9, Myh10/MYH10 and Myh14/MYH14 [25]. Distinct paralogs co-assemble into the same myosin mini-filaments [26] and to some extent seem to have redundant actions within the cells. However, several in vitro studies point to specific roles of NMHC paralogs. For example, Myh9 plays a key role in setting the speed of furrow ingression during cytokinesis [27,28] and is essential to drive bleb retraction [29]. During cell-cell contact formation, Myh9 was found essential for cadherin adhesion molecule clustering and setting contact size while Myh10 would be involved in force transmissions across the junction and would influence contact rearrangements [30,31].

These studies at the subcellular level and short timescale complement those at the organismal level and long timescale. In mice, zygotic knockout of Myh14 causes no obvious phenotype with animals surviving to adulthood with no apparent defect [32] whereas the loss of either Myh9 [25] or Myh10 [33] is embryonic lethal. Myh9 zygotic knockout embryos die at E7.5 due to visceral endoderm adhesion defects [25]. Myh10 zygotic knockout mice die between E14.5 and P1 because of heart, brain and liver defects [33]. In addition, knocking out both Myh10 and Myh14 can lead to abnormal cytokinesis [32]. Elegant gene replacement experiments also revealed insights into partial functional redundancy between Myh9 and Myh10 during development [34]. However, despite the prominent role of actomyosin contractility during preimplantation development [32], the specific functions of NMHC paralogs remain largely unknown. Previous genetic studies relied on zygotic knockouts [25,32,33], which do not remove the maternally deposited mRNA and proteins of the deleted genes. This often hides the essential functions of genes during preimplantation morphogenesis, as is the case, for example, with the cell-cell adhesion molecule Cdh1 [35]. Moreover, NMHCs could have redundant functions and gene deletions may trigger compensation mechanisms, which obscure the function of essential genes [36].

In this study, we generate maternal-zygotic deletions of single or double NMHC genes to investigate the molecular control of contractile forces during preimplantation development. We use nested time-lapse microscopy to quantitatively assess the effect of maternal-zygotic deletions at different timescales. This reveals the dominant role of Myh9 over Myh10 in generating the contractility which shapes the mouse blastocyst. In addition, double maternal-zygotic and Myh10 knockout compensatory Myh9 reveals mechanisms provided by Myh10 in generating enough contractility for cytokinesis when Myh9 is absent. Moreover, maternal-zygotic knockout of both Myh9 and Myh10 can cause embryos to fail all five successive cleavages, resulting in syncytial single-celled embryos. These single-celled embryos nevertheless initiate lineage specification and blastocoel formation by accumulating fluid into intracellular vacuoles. Therefore, double maternal-zygotic NMHC mutants reveal that fluid accumulation in the blastocyst is a cell-autonomous process. Finally, we confirm this surprising finding by fusing all blastomeres of WT embryos, thereby forming single-celled embryos, which accumulate fluid into inflating vacuoles.

## Results

### NMHC paralogs during preimplantation development

As in human, the mouse genome contains 3 genes encoding NMHCs: Myh9, Myh10 and Myh14. To decipher the specific contributions of NMHC paralogs to preimplantation development, we first investigated the expression of Myh9, Myh10 and Myh14.

We performed real time quantitative PCR (qPCR) at 4 different stages in order to cover the levels of transcripts at key steps of preimplantation development. We detect high levels of Myh9 mRNA throughout preimplantation development (Fig S1A). Importantly, transcripts of Myh9 are by far the most abundant among NMHCs at the zygote stage (E0.5), which suggests that Myh9 is the main NMHC paralog provided maternally. Myh10 mRNA is detected at very low levels in zygotes before it reaches comparable levels to Myh9 mRNA at the morula stage (Fig S1A). We find that Myh14 is not expressed during preimplantation stages (Fig S1A). Since Myh14 homozygous mutant mice are viable and show no apparent defects [32], Myh14 is unlikely to play an important role during preimplantation development. These qPCR measurements are in agreement with available mouse single cell RNA sequencing (scRNA-seq) data (Fig S1B) [37]. In human, scRNA-seq data indicate similar expression levels between MYH9 and MYH10 during preimplantation development and, as in mouse, the absence of MYH14 expression (Fig S1C) [38]. This points to a conserved regulation of NMHC paralogs in mouse and human preimplantation development.

At the protein level, immunostaining of Myh9 becomes visible at the cortex of blastomeres from the zygote stage onwards (Fig S1D). On the other hand, Myh10 becomes detectable at the earliest at the 16-cell stage (Fig S1D). Finally, we used transgenic mice expressing endogenously tagged Myh9-GFP to assess the relative parental contributions of Myh9 protein (Fig S1E-F). Embryos coming from Myh9-GFP females show highest levels of fluorescence at the zygote stage, consistent with our qPCR measurement and scRNA-seq data (Fig S1A-B). Myh9-GFP produced from the paternal allele is detected at the 4-cell stage and increases until blastocyst stage, reaching levels comparable to blastocyst coming from Myh9-GFP females (Fig S1E-F).

Together, we conclude that Myh9/MYH9 and Myh10/MYH10 are the most abundant NMHCs during mouse and human preimplantation development and that Myh9 is heavily maternally provided at both the transcript and protein level in the mouse embryo (Fig S1).

### Preimplantation development of single maternal-zygotic knockouts of Myh9 or Myh10

Initial studies of Myh9 or Myh10 zygotic knockouts reported that single zygotic knockouts are able to implant, suggesting they are able to form a functional blastocyst [25,33]. A potential lack of phenotype could be due to maternally provided products, which are most abundant in the case of Myh9 (Fig S1). To eliminate this contribution, we used Zp3::Cre mediated maternal deletion of conditional knockout alleles of Myh9 and Myh10. This generated either maternal-zygotic (mz) homozygous or maternal only (m) heterozygous knockout embryos for either Myh9 or Myh10. Embryos were recovered at E1.5, imaged throughout the rest of their preimplantation development, stained and genotyped once wild-type (WT) embryos reached the blastocyst stage. We implemented a nested time-lapse protocol to image each embryo on the long (every 30 min for about 50 h) and short (every 5 s for 10 min twice for each embryo around the time of their 8-cell stage) timescales. This imaging protocol allowed us to visualize the effect of actomyosin contractility at multiple timescales as it is involved in pulsatile contractions, cytokinesis and morphogenesis, which take place on timescales of tens of seconds, minutes and hours, respectively [8,13].

The first visible phenotype concerns the zona pellucida (ZP), which encapsulate the preimplantation embryo. WT and Myh10 mutants have a spherical ZP whereas in Myh9 mutants show an irregularly shaped ZP (Fig S2A-B). This suggests a previously unreported role of contractility in the formation of the ZP during oogenesis that is mediated specifically by maternal Myh9. The abnormally shaped ZP could cause long term deformation of the embryo as previous studies indicate that the ZP can influence the shape of the embryo at the blastocyst stage [39,40].

During compaction, the first morphogenetic movement of the mouse embryo, contact angles increase from 87 ± 3° to 147 ± 2° between the 3^rd^ and 4^th^ cleavage in WT embryos (mean ± SEM, 23 embryos, Fig 1A-B, Table 1-2, Movie 1), as measured previously [13,14]. In mzMyh9−/− embryos contact angles only grow from 85 ± 2° to 125 ± 4° (mean ± SEM, 15 embryos, Fig 1A-B, Table 1-2). This reduced ability to compact is in agreement with previous measurements on heterozygous mMyh9+/− embryos, which generate a lower surface tension and therefore cannot efficiently pull cells into closer contact [8]. During the 8-cell stage, mzMyh10−/− embryos initially grow their contacts from 87 ± 5° to 121 ± 4°, similarly to mzMyh9−/− embryos (mean ± SEM, 11 embryos, Fig 1A-B, Table 1). However, unlike mzMyh9−/− embryos, mzMyh10−/− embryos eventually reach compaction levels identical to WT embryos by the end of the 16-cell stage (148 ± 4°, mean ± SEM, 11 embryos, Fig 1A-B, Table 1). Heterozygous mMyh9+/− or mMyh10+/− embryos show similar phenotypes to their respective homozygous counterparts (Fig S3A-B, Table 1-2), suggesting that maternal loss dominates for both NMHC paralogs. Together, we conclude that Myh9 is essential for embryos to compact fully whereas Myh10 only regulates the rate of compaction.

**Table 1:** Statistical analysis of compaction data. Mean contact angles and developmental times related to Fig 1B, 3B, S4A and S6A with associated SEM, embryo number and p value from unpaired two-tailed Welch’s t-testing against WT.

|  |  | Time after 3rd cleavage |  |  |  | Time before initial 4th cleavage |  |  |  | Time after 4th cleavage |  |  |  | Time before initial 5th cleavage |  |  |  |
| --- | --- | --- | --- | --- | --- | --- | --- | --- | --- | --- | --- | --- | --- | --- | --- | --- | --- |
|  |  | mean | SEM | embryos | p value | mean | SEM | embryos | p value | mean | SEM | embryos | p value | mean | SEM | embryos | p value |
| WT | Time after 3rd cleavage [h] | 0 | 0 | 23 |  | 7.0 | 0.3 | 23 |  | 10.9 | 0.3 | 21 |  | 16.7 | 0.4 | 22 |  |
|  | Contact angle [°] | 87 | 3 |  |  | 147 | 2 |  |  | 140 | 3 |  |  | 148 | 2 |  |  |
| mzMyh9<sup>-/-</sup> | Time after 3rd cleavage [h] | 0 | 0 | 15 |  | 9.8 | 0.5 | 15 | 0.0001 | 13.8 | 0.6 |  | 0.002 | 18.8 | 0.5 |  | 0.01 |
|  | Contact angle [°] | 85 | 2 |  | 0.5 | 125 | 4 |  | 0.00001 | 104 | 5 | 10 | 0.00001 | 131 | 8 | 8 | 0.02 |
| mMyh9<sup>+/-</sup> | Time after 3rd cleavage [h] | 0 | 0 | 8 |  | 10.4 | 0.7 | 8 | 0.001 | 14.6 | 0.8 |  | 0.002 | 18.7 | 0.7 |  | 0.002 |
|  | Contact angle [°] | 86 | 5 |  | 0.5 | 115 | 6 |  | 0.0001 | 112 | 7 | 8 | 0.003 | 113 | 21 | 3 | 0.1 |
| mzMyh10<sup>-/-</sup> | Time after 3rd cleavage [h] | 0 | 0 | 11 |  | 5.4 | 0.4 | 11 | 0.004 | 10.2 | 0.6 |  | 0.2 | 15.8 | 0.4 |  | 0.1 |
|  | Contact angle [°] | 87 | 5 |  | 0.8 | 121 | 4 |  | 0.0001 | 126 | 5 | 11 | 0.03 | 148 | 4 | 11 | 0.8 |
| mMyh10<sup>+/-</sup> | Time after 3rd cleavage [h] | 0 | 0 | 20 |  | 6.3 | 0.4 |  | 0.2 | 10.8 | 0.3 |  | 0.5 | 16.8 | 0.6 |  | 0.9 |
|  | Contact angle [°] | 85 | 3 |  | 0.3 | 128 | 5 | 20 | 0.0004 | 128 | 3 | 20 | 0.006 | 149 | 2.8 | 20 | 0.5 |
| mzMyh9<sup>-/-</sup>;mzMyh10<sup>-/-</sup> | Time after 3rd cleavage [h] | 0 | 0 | 7 |  | 9.7 | 0.8 |  | 0.07 | 12 |  | 1 |  |  |  |  |  |
|  | Contact angle [°] | 89 | 6 |  | 0.8 | 117 | 4 | 3 | 0.006 | 84.5 |  |  |  |  |  |  |  |
| mzMyh9<sup>-/-</sup>;mMyh10<sup>+/-</sup> | Time after 3rd cleavage [h] | 0 | 0 | 7 |  | 6.9 | 1.7 | 4 | 1.0 | 9 | 0.5 | 2 | 0.1 | 12 |  | 1 |  |
|  | Contact angle [°] | 83 | 5 |  | 0.6 | 100 | 5 |  | 0.002 | 108 | 14 |  | 0.07 | 113 |  |  |  |
| mMyh9<sup>+/-</sup>;mMyh10<sup>+/-</sup> | Time after 3rd cleavage [h] | 0 | 0 | 7 |  | 8 | 1.3 | 5 | 0.5 | 11 | 0.5 | 3 | 1.0 |  |  |  |  |
|  | Contact angle [°] | 76 | 7 |  | 0.1 | 110 | 5 |  | 0.0006 | 105 | 3 |  | 0.002 |  |  |  |  |

**Figure 1:**
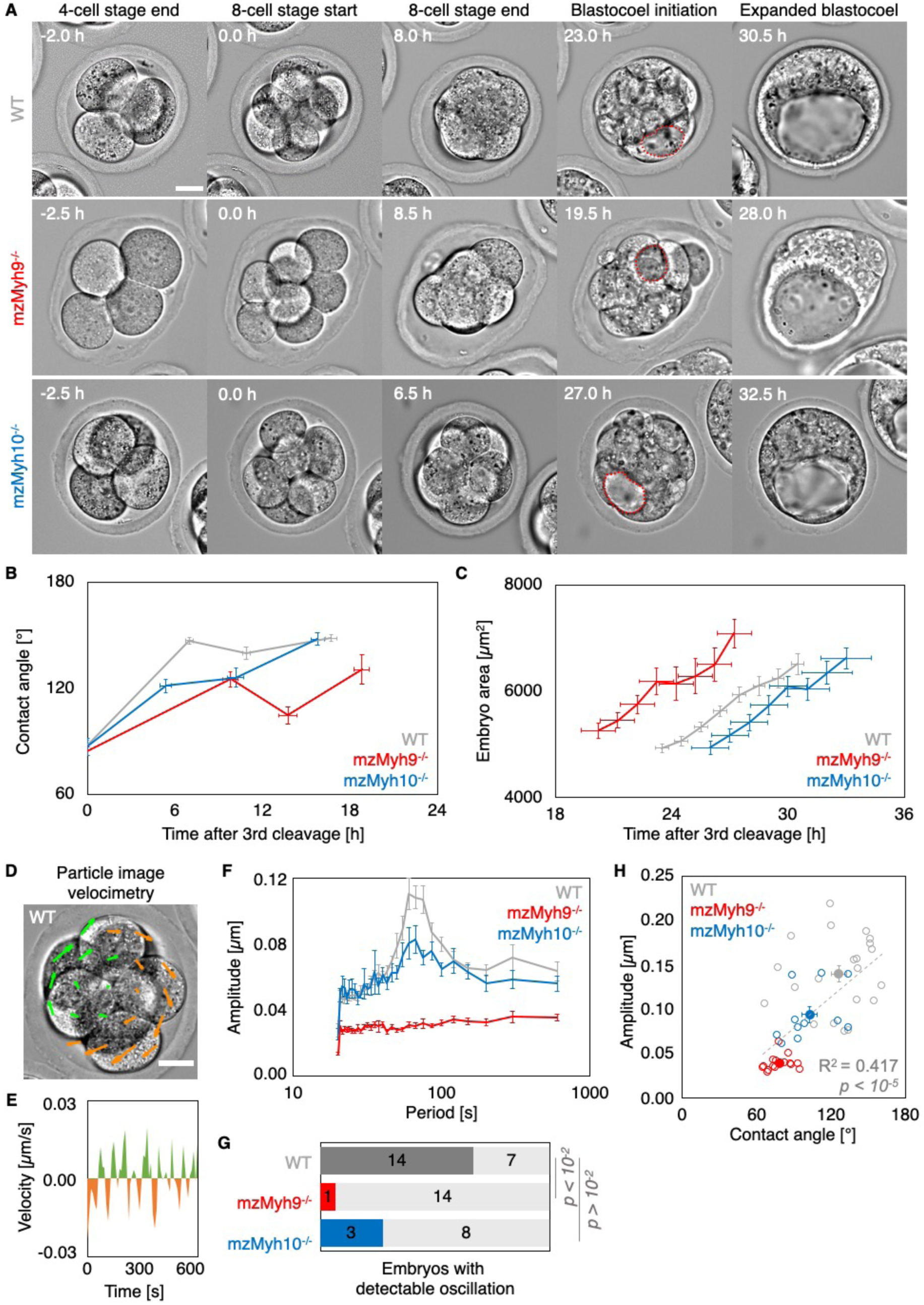
Multiscale analysis of morphogenesis in single maternal-zygotic Myh9 or Myh10 mutant embryos. A) Representative images of long term time-lapse of WT, mzMyh9−/− and mzMyh10−/− embryos at the end of the 4-cell stage, start and end of the 8-cell stage, at the initiation of blastocoel formation and early blastocyst stage (Movie 1). Scale bar, 20 µm. Time in hours after 3^rd^ cleavage. Dotted red lines indicate the nascent lumen. B) Contact angle of WT (grey, n = 23, 23, 21, 22), mzMyh9−/− (red, n = 15, 15, 10, 8) and mzMyh10−/− (blue, n = 11, 11, 11, 11) embryos after the 3^rd^ cleavage, before and after the 4^th^ cleavage and before the 5^th^ cleavage. Data show mean ± SEM. Statistical analyses are provided in Table 1. C) Embryo growth during lumen formation for WT (grey, n = 20), mzMyh9−/− (red, n = 9) and mzMyh10−/− (blue, n = 7) embryos measured for 7 continuous hours after a lumen of at least 20 µm is observed (as in blastocoel initiation of panel A). Data show mean ± SEM. D) Representative image of short term time-lapse overlaid with a subset of velocity vectors from Particle Image Velocimetry (PIV) analysis. Green for positive and orange for negative Y-directed movement. E) Velocity over time for a representative velocity vector of embryo shown in D and Movie 2. F) Power spectrum resulting from Fourier transform of PIV analysis of WT (grey, n = 21), mzMyh9−/− (red, n = 15) and mzMyh10−/− (blue, n = 11) embryos. Peaks indicate periodic movements at a given period (see methods). Data show mean± SEM. G) Proportion of WT (grey, n = 21), mzMyh9−/− (red, n = 15) and mzMyh10−/− (blue, n = 11) embryos showing detectable oscillations in their power spectrum (see methods). Chi^2^ test p value comparing to WT is indicated. H) Amplitude of oscillation as a function of the contact angle for WT (grey, n = 21), mzMyh9−/− (red, n = 15) and mzMyh10−/− (blue, n = 11) embryos. Open circle show individual embryos and filled circles give mean ± SEM of a given genotype. Pearson’s R^2^ and p value are indicated. Statistical analyses are provided in Table 3.

During the 8-cell stage, contractility becomes visible on the short timescale via the presence of periodic contractions, which we can use to gauge the specific contribution of NMHCs [8,13]. We performed particle image velocimetry (PIV) and Fourier analysis to evaluate the period and amplitude of periodic movements (Fig 1D-F) [13]. If 14/21 WT embryos displayed periodic contractions, these were rarely detected in Myh9 mutants (1/15 mzMyh9−/− and 0/8 mMyh9+/− embryos) and occasionally in Myh10 mutants (3/11 mzMyh10−/− and 9/21 mMyh10+/− embryos, Fig 1G and Fig S3D, Movie 2). This suggests that contractility is reduced after maternal loss of Myh10 and greatly reduced following maternal loss of Myh9. This hierarchy in the phenotypes of the NMHC paralog mutants parallels the one observed on the long timescale during compaction (Fig 1B). In fact, we find that the amplitude of periodic contractions correlates with the contact angle across the genotypes we considered (Fig 1H and S3E, 78 embryos, Pearson’s *R* = 0.636, *p* < 10^−5^, Table 3). This analysis across timescales reveals the continuum between the short term impact of Myh9 or Myh10 onto contractile movements and the long term morphogenesis, as previously observed for internalizing ICM cells [8].

**Table 2:** Statistical analysis of cell cycle duration data. Mean durations of 8-cell stage, 4^th^ wave of cleavages and 16-cell stage related to Fig 1B, 3B, S4A and S6A with associated SEM, embryo number and p value from unpaired two-tailed Welch’s t-testing against WT.

|  | Duration of 8-cell stage [h] |  |  |  | Duration of 4th wave of cleavages [h] |  |  |  | Duration of 16-cell stage [h] |  |  |  |
| --- | --- | --- | --- | --- | --- | --- | --- | --- | --- | --- | --- | --- |
|  | mean | SEM | embryos | p value | mean | SEM | embryos | p value | mean | SEM | embryos | p value |
| WT | 7.0 | 0.3 | 23 |  | 4.0 | 0.3 | 23 |  | 5.5 | 0.4 | 23 |  |
| mzMyh9<sup>-/-</sup> | 9.8 | 0.5 | 15 | 0.0001 | 4.8 | 0.7 | 15 | 0.3 | 5.7 | 0.8 | 14 | 0.8 |
| mMyh9<sup>+/-</sup> | 10.4 | 0.7 | 8 | 0.001 | 4.2 | 1.2 | 8 | 0.9 | 7.0 | 1.1 | 8 | 0.1 |
| mzMyh10<sup>-/-</sup> | 5.4 | 0.4 | 11 | 0.004 | 4.8 | 0.8 | 11 | 0.2 | 5.6 | 0.5 | 11 | 0.9 |
| mMyh10<sup>+/-</sup> | 6.3 | 0.4 | 20 | 0.2 | 4.5 | 0.4 | 20 | 0.3 | 6.1 | 0.5 | 20 | 0.4 |
| mzMyh9<sup>-/-</sup>;mzMyh10<sup>-/-</sup> | 10.3 | 1.4 | 8 | 0.06 | 2.0 | 0.8 | 3 | 0.1 |  |  |  |  |
| mzMyh9<sup>-/-</sup>;mMyh10<sup>+/-</sup> | 8.1 | 1.4 | 6 | 0.5 | 4.3 | 1.1 | 3 | 0.8 |  |  |  |  |
| mMyh9<sup>+/-</sup>;mMyh10<sup>+/-</sup> | 9.3 | 1.0 | 8 | 0.06 | 3.6 | 0.7 | 5 | 0.6 |  |  |  |  |

**Table 3:** Statistical analysis of periodic contraction data. Mean amplitude of periodic movements and contact angle related to Fig 1H and S3E with associated SEM, embryo number and p value from unpaired two-tailed Welch’s t-testing against WT.

|  | embryos | Maximum amplitude within 50-120 s oscillation period range [μm] |  |  | Contact angle [°] |  |  |
| --- | --- | --- | --- | --- | --- | --- | --- |
|  |  | mean | SEM | p value | mean | SEM | p value |
| WT | 21 | 0.140 | 0.009 |  | 126 | 5 |  |
| mzMyh9<sup>-/-</sup> | 15 | 0.039 | 0.002 | 0.00003 | 78 | 3 | 0.000003 |
| mMyh9<sup>+/-</sup> | 8 | 0.045 | 0.003 | 0.00005 | 82 | 6 | 0.001 |
| mzMyh10<sup>-/-</sup> | 11 | 0.094 | 0.009 | 0.01 | 103 | 6 | 0.02 |
| mMyh10<sup>+/-</sup> | 21 | 0.111 | 0.013 | 0.01 | 108 | 5 | 0.02 |

We also noted that the duration of the 8-cell stage is longer in embryos lacking maternal Myh9 (9.8 ± 0.5 h from the 3^rd^ to the 4^th^ wave of cleavages, mean ± SEM, 15 embryos) as compared to WT (7.0 ± 0.3 h, mean ± SEM, 23 embryos, Fig 1A-B, Table 1-2, Movie 1). This is not the case for the duration of the 4^th^ wave of cleavages or the ensuing 16-cell stage, which are similar in these different genotypes (Table 2). On the long timescale, longer cell cycles could affect the number of cells in Myh9 mutants. Indeed, when reaching the blastocyst stage, mzMyh9−/− embryos count less than half the number of cells than WT (58.1 ± 2.9 cells in 23 WT embryos, as compared to 25.2 ± 2.8 cells in 15 mzMyh9−/− embryos, Mean ± SEM, Fig 2B). Considering individual mzMyh9−/− embryos, the number of cells at blastocyst stage is best explained by the duration of the 8-cell stage (15 embryos, Pearson’s *R* = −0.610, *p* < 10^−2^) than by the duration of the 4^th^ cleavage wave (15 embryos, Pearson’s *R* = 0.133, *p* >10^−2^) or of the 16 cell-stage (14 embryos, Pearson’s *R* = −0.071, *p* > 10^−2^). Therefore, the lengthened 8-cell stage of mzMyh9−/−may be a primary cause for their reduced cell number. Reduced cell number could also result from the occasional cytokinesis defects observed in maternal Myh9 mutants such as cleavages that reverse back shortly after (Movie 3). Loss of Myh9 is likely to cause difficulties during cytokinesis [27,28], which could in turn impact cell cycle progression (Fig 1B) and explain the significant reduction in cell number in mzMyh9−/− embryos. On the other hand, mzMyh10−/− embryos do not show cell cycle delay and make blastocysts with the correct cell number (Fig 1B and 2B), indicating that Myh9 is the primary NMHC powering cytokinesis during mouse preimplantation development.

**Figure 2:**
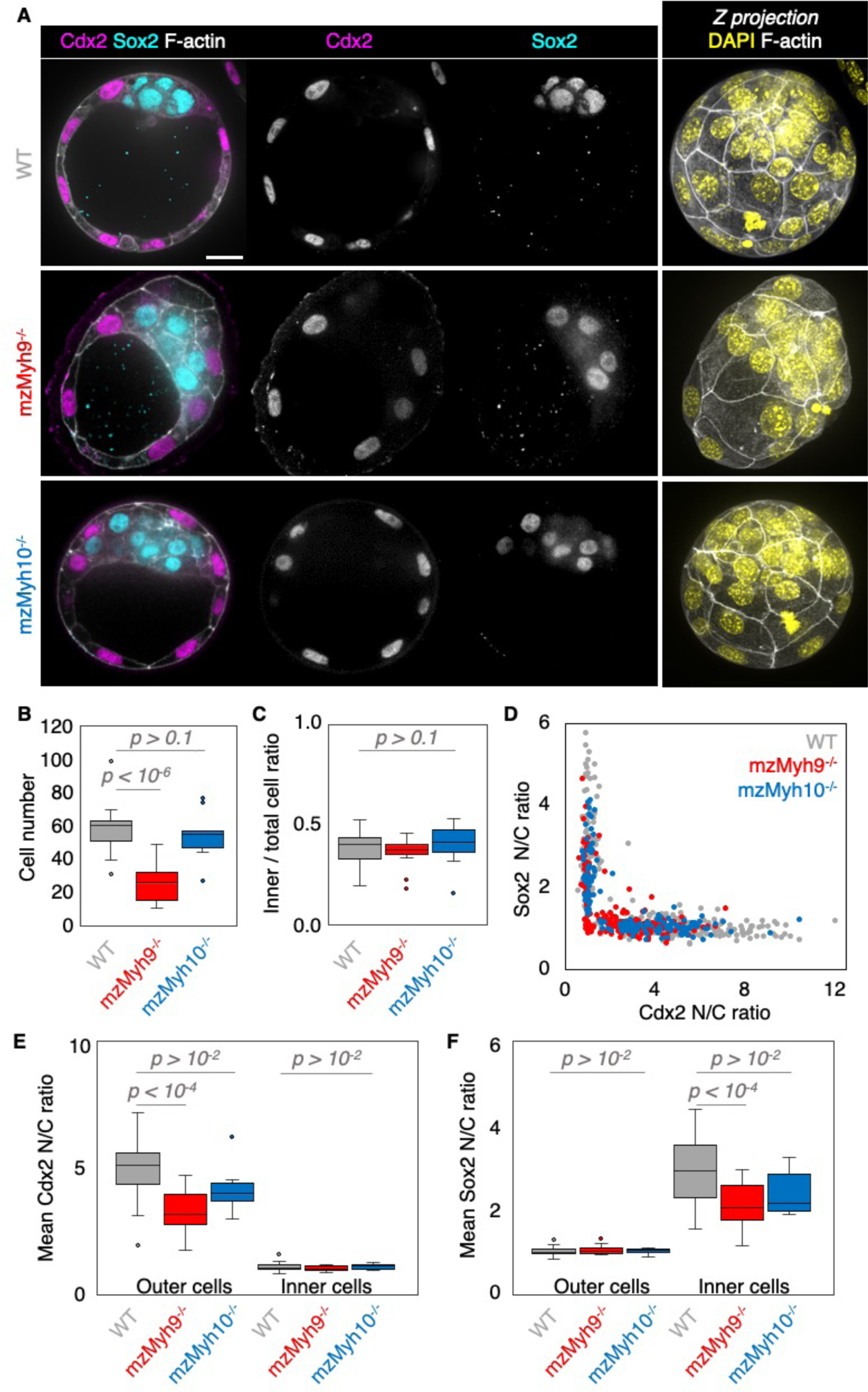
TE and ICM lineages analysis of single maternal-zygotic Myh9 or Myh10 mutant embryos. A) Representative images of WT, mzMyh9−/− and mzMyh10−/− embryos stained for TE and ICM markers Cdx2 (magenta) and Sox2 (cyan). DAPI in yellow and F-actin in grey. The same mutant embryos as in Fig 1A are shown. B-C) Total cell number (B) and proportion of inner cells (C) in WT (grey, n = 23), mzMyh9−/− (red, n = 15) and mzMyh10−/− (blue, n = 11) embryos. D) Nuclear to cytoplasmic (N/C) ratio of Cdx2 and Sox2 staining for individual cells from WT (grey, n = 345), mzMyh9−/− (red, n = 204) and mzMyh10−/− (blue, n = 160) embryos. E-F) N/C ratio of Cdx2 (E) and Sox2 (F) staining for outer (left) or inner (right) cells from averaged WT (grey, n = 23), mzMyh9−/− (red, n = 15) and mzMyh10−/− (blue, n = 11) embryos. Mann-Whitney U test p values comparing to WT are indicated.

The 2^nd^ morphogenetic movement consists in the positioning of cells on the inside of the embryo after the 4^th^ cleavage division to form the first lineages of the mammalian embryo. To see if this process is affected in NMHC mutants, we counted the number of inner and outer cells on immunostaining at the blastocyst stage (Fig 2A-C). Despite showing less than half the expected cell number, mzMyh9−/− blastocysts show the same proportion of inner and outer cells as WT and mzMyh10−/− embryos (Fig 2C). This suggests that the remaining contractility is sufficient to drive cell internalization or that oriented cell divisions can mitigate the loss of contractility-mediated internalization [6–8]. To assess whether differentiation itself is affected, we performed immunostaining of Cdx2 as a TE marker and Sox2 as an ICM marker (Fig 2A). mzMyh10−/− embryos display negligible reduction in Cdx2 and Sox2 levels compared to WT embryos (Fig 2D-F). On the other hand, we find that mzMyh9−/− embryos show lower levels of Cdx2 in their outer cells, as measured here and previously for mMyh9+/− embryos (Fig 2D-E and Fig S4) [8], and of Sox2 for inner cells compared to WT embryos (Fig 2F). This is caused in part by the presence of individual unspecified cells localized both inside and at the surface of mzMyh9−/− embryos (in Fig 2D, cells with nuclear to cytoplasmic ratios of ∼1 for both Cdx2 and Sox2 belong to both inner and outer cells from multiple embryos). To assess whether delayed cell cycle progression may explain the reduced differentiation, we calculated the correlation between the duration of the 8-cell stage and the levels of Cdx2 in the TE and of Sox2 in the ICM. We find no significant correlation among mzMyh9−/−embryos (Pearson’s *R* = −0.325 for Cdx2 and −0.472 for Sox2, 15 embryos, *p* > 10^−2^) or when extending to all maternal Myh9 mutants (Pearson’s *R* = −0.193 for Cdx2 and −0.076 for Sox2, 22 embryos, *p* > 10^−2^). This does not support reduced differentiation levels due to cell cycle delay. Together, these findings are consistent with Myh9-mediated contractility being required for the correct expression of lineage markers within the blastocyst beyond cell positioning [8].

The final morphogenetic step shaping the blastocyst is the formation of the first mammalian lumen. After the 5^th^ cleavage, the growth of the blastocoel inflates and increases the projected area of WT embryos steadily at 3.9 ± 0.5 µm^2^/min (mean ± SEM, 20 embryos, Fig 1C, Movie 1). During lumen formation, mzMyh9−/− and mzMyh10−/− embryos inflate at 3.9 ± 1.0 and 4.0 ± 0.5 µm^2^/min respectively, which is similar to WT embryos (mean ± SEM, 9 and 7 embryos, Student’s *t* test comparing to WT *p* > 10^−2^, Fig 1C). This suggests that blastocoel expansion rate is unaffected in NMHC mutants. Importantly, we observe that the time of lumen opening in mzMyh9−/− embryos is not delayed compared to WT embryos (Fig 1A-C). In fact, the lumen inflates on average at an earlier stage in mzMyh9−/− embryos than in WT (Fig 1C). Indeed, some mzMyh9−/− embryos begin inflating their lumen before the 5^th^ wave of cleavages (5/11, Movie 1), while all 20 WT and 7 mzMyh10−/− embryos begin lumen formation after the first cleavages of the 5^th^ wave. This argues against mzMyh9−/− embryos having a global delay in their development, despite their longer cell cycles, lower cell number (Fig 1B and Fig 2B) and impaired differentiation (Fig 2D-F). Finally, when the lumen becomes sufficiently large, embryos come into contact with the ZP. 3D segmentation of blastocysts reveals that embryos with mutated maternal Myh9 alleles become less spherical than those with a WT allele of Myh9 (Fig S2C-F), which suggests that the misshapen ZP could deform the embryo. This is consistent with previous studies reporting on Myh9 mutant embryos being more deformable than WT [8] and on the influence of the ZP on the shape of the blastocyst [39,40].

Maternal Myh9 mutants show defective cytokinesis and reduced cell number, which could constitute the cause for the defects in morphogenesis and lineage specification. To separate the effects of impaired contractility and of defective cell divisions, we decided to reduce cell number by fusing blastomeres of WT embryos together. Reducing cell number by ¼ or ½ at the 4-cell stage results in embryos compacting with the same magnitude and forming a lumen concomitantly to control embryos (Fig S5A-C, Movie 4). Also, embryos with reduced cell number differentiate into TE and ICM identically to control embryos as far as lineage proportions and fate marker levels are concerned (Fig S5D-H). Therefore, the blastocyst morphogenesis and lineage specification defects observed in maternal Myh9 mutants are not likely to result simply from cytokinesis defects and their impact on cell number.

Together, these analyses reveal the critical role of Myh9 in setting global cell contractility on short and long timescales in order to effectively drive compaction, cytokinesis and lineage specification. Comparably, the function of Myh10 during early blastocyst morphogenesis is less prominent. Importantly, the similarity of maternal homozygous and maternal heterozygous mutants indicate that the preimplantation embryo primarily relies on the maternal pools of Myh9. We conclude that maternally provided Myh9 is the main NMHC powering actomyosin contractility during early preimplantation development.

### Preimplantation development of double maternal-zygotic Myh9 Myh10 knockout

Despite Myh9 being the main NMHC provided maternally, the successful cleavages, shape changes and differentiation observed in mMyh9+/− and mzMyh9−/− embryos suggest that some compensation by Myh10 could occur in these embryos. To test for compensations between the two NMHC paralogs expressed during development, we preimplantation generated double maternal-zygotic knockout embryos. Myh9 and Myh10 (mzMyh9−/−;mzMyh10−/−) embryos.

Nested time-lapse revealed that mzMyh9−/−;mzMyh10−/− embryos fail most attempts of cytokinesis (Movie 5). In addition to failed cytokinesis, mzMyh9−/−;mzMyh10−/− embryos show increased cell cycle durations, which are more severe than for mzMyh9−/− embryos (Fig 3A-B, Table 1-2). This results in mzMyh9−/−;mzMyh10−/− embryos with only 2.9 ± 0.5 cells when reaching the blastocyst stage (Fig 4B). In fact, they occasionally develop into blastocyst-stage embryos consisting of only one single cell (1/8 embryo). Of those which succeed in dividing at least once, compaction after the 3^rd^ cleavages is weak with contact angles for double mutants at capping at 117 ± 4° compared to 147 ± 2° for WT (mean ± SEM, 3 and 23 embryos, Fig 3B, Table 1, Movie 5). Consistently with previous observations, periodic contractions are undetectable in any of the double mutants that we analyzed (Fig 3D-E and Fig S6C-D, Movie 2). Together, this indicates that double maternal-zygotic Myh9 Myh10 knockout embryos generate extremely weak contractile forces compared to WT, single maternal-zygotic Myh9 or Myh10 knockout embryos.

**Figure 3:**
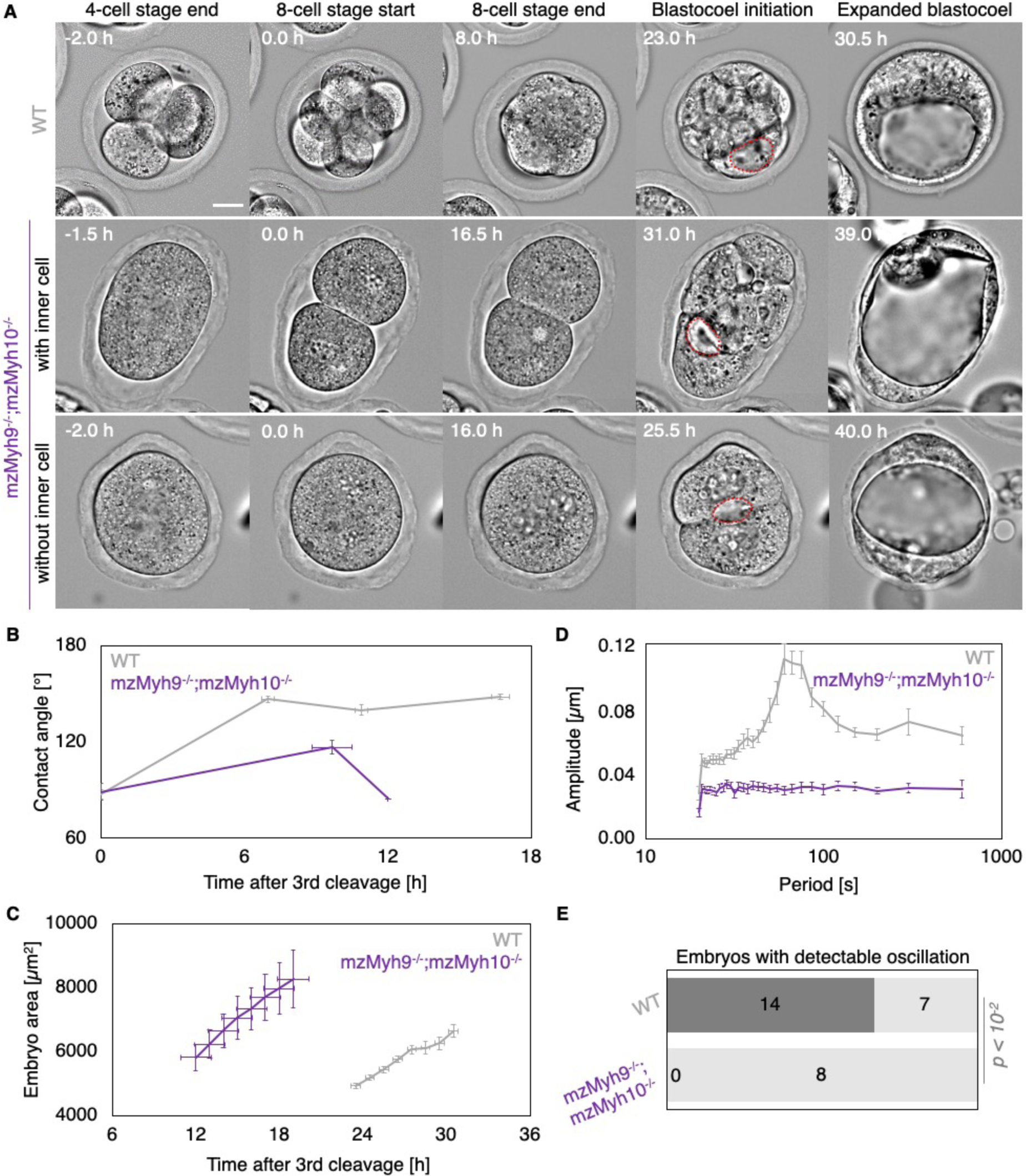
Multiscale analysis of morphogenesis in double maternal-zygotic Myh9 and Myh10 mutant embryos. A) Representative images of long term time-lapse of WT and mzMyh9−/−;mzMyh10−/− embryos at the end of the 4-cell stage, start and end of the 8-cell stage, at the initiation of blastocoel formation and early blastocyst stage (Movie 5). Middle row shows an embryo that cleaved once at the time of the 3^rd^ cleavage and twice during the 4^th^ cleavage wave. Bottom row shows an embryo that cleaved once during at the time of the 4^th^ cleavage. B) Contact angle of WT (grey, n = 23, 23, 21, 22) and mzMyh9−/−;mzMyh10−/− (purple, n = 7, 3, 1) embryos after the 3^rd^ cleavage, before and after the 4^th^ cleavage and before the 5^th^ cleavage. Data show mean ± SEM. Statistical analyses are provided in Table 1-2. C) Embryo growth during lumen formation for WT (grey, n = 20) and mzMyh9−/−;mzMyh10−/− (purple, n = 5) embryos measured for 7 continuous hours after a lumen of at least 20 µm is observed (as in blastocoel initiation of panel A). Data show mean ± SEM. D) Power spectrum resulting from Fourier transform of PIV analysis of WT (grey, n = 21) and mzMyh9−/−;mzMyh10−/− (purple, n = 8) embryos (Movie 2). Data show mean ± SEM. F) Proportion of WT (grey, n = 21) and mzMyh9−/−;mzMyh10−/− (purple, n = 8) embryos showing detectable oscillations in their power spectrum. Chi^2^ test p value comparing to WT is indicated.

**Figure 4:**
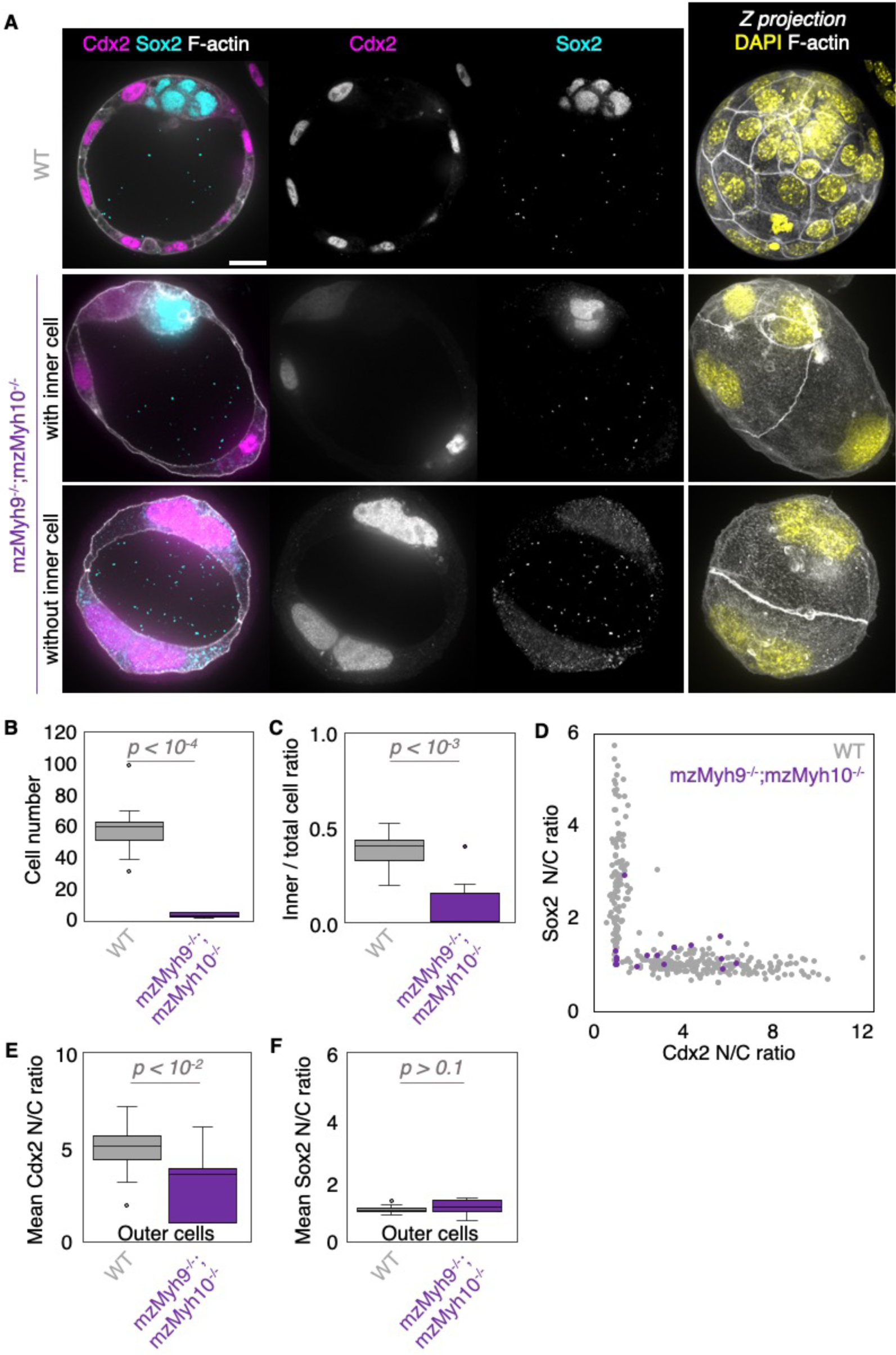
TE and ICM lineages analysis of double maternal-zygotic Myh9 and My10 mutant embryos. A) Representative images of WT and mzMyh9−/−;mzMyh10−/− embryos stained for TE and ICM markers Cdx2 (magenta) and Sox2 (cyan). DAPI in yellow and F-actin in grey. The same mutant embryos as in Fig 3A are shown. B-C) Total cell number (B) and proportion of inner cells (C) in WT (grey, n = 23) and mzMyh9−/−;mzMyh10−/− (purple, n = 8) embryos. D) Nuclear to cytoplasmic (N/C) ratio of Cdx2 and Sox2 staining for individual cells from WT (grey, n = 345) and mzMyh9−/−;mzMyh10−/− (purple, n = 17) embryos. E-F) N/C ratio of Cdx2 (E) and Sox2 (F) staining for outer cells from averaged WT (grey, n = 23) and mzMyh9−/−;mzMyh10−/− (purple, n = 7) embryos. Mann-Whitney U test p values comparing to WT are indicated.

When investigating fate specification, Cdx2 is present in 5/7 embryos as compared to all 23 WT embryos we analyzed (Fig 4E). Sox2 is only found when embryos succeed in internalizing at least one cell (Fig 4A and D). The levels of Cdx2 in outer cells are reduced compared to WT embryos, as observed previously for single maternal-zygotic Myh9 knockout embryos. Nevertheless, these measurements suggest that the lineage specification program is still being executed in mzMyh9−/−;mzMyh10−/− embryos despite the absence of observable contractility.

The presence of Cdx2 suggests that mzMyh9−/−;mzMyh10−/− embryos can initiate differentiation into TE, the epithelium responsible for making the blastocoel. The ability of mzMyh9−/−;mzMyh10−/− embryos to create *de novo* a functional polarized epithelium is further supported by the fact that they can eventually proceed with polarized fluid accumulation and create a blastocoel (Fig 3A, Movie 5). As observed in single maternal Myh9 mutants, despite extended cell cycles, mzMyh9−/−;mzMyh10−/− embryos are not delayed to initiate lumen formation (Fig 3C, Movie 5). Also, we measure that mutant embryos grow at rates that are similar to WT ones (5.8 ± 1.5 and 3.9 ± 0.5 µm^2^/min, mean ± SEM, 5 and 20 embryos, Student’s *t* test *p* > 10^−2^, Fig 3C). This indicates that contractility does not influence the initial growth rate of the blastocoel and that blastocoel formation does not require powerful actomyosin contractility.

Finally, we measure similar metrics for the single and double heterozygous Myh9 and Myh10 mutant embryos (Fig S6-7, Table 1), indicating once again that maternal contributions predominantly set the contractility for preimplantation development. We conclude that after maternal loss of both Myh9 and Myh10, contractility is almost entirely abolished. This effect is much stronger than after the loss of maternal Myh9 only and suggests that, in single maternal Myh9 mutants, Myh10 can compensate significantly, which we would not anticipate from our previous single mzMyh10−/− knockout analysis. Therefore, despite Myh9 being the main engine of preimplantation actomyosin contractility, Myh10 ensures a substantial part of blastocyst morphogenesis in the absence of Myh9. The nature and extent of this compensation will need further characterization.

### Preimplantation development of single-celled embryos

Embryos without maternal Myh9 and Myh10 can fail all successive cleavages when reaching the time of the blastocyst stage (6/25 embryos all maternal Myh9 and Myh10 mutant genotypes combined, Fig 5A, Movie 6). In single-celled embryos, multiple cleavage attempts at intervals of 10-20 h can be observed, suggesting that the cell cycle is still operational in those embryos (Movie 5). This is further supported by the presence of multiple large nuclei in single-celled embryos (Fig 5C). Interestingly, these giant nuclei contain Cdx2, and no Sox2, suggesting that, in addition to the cell cycle, the lineage specification program is still operational (Fig 5C-E). Strikingly, we also observed that at the time of blastocoel formation, single-celled maternal Myh9 and Myh10 mutant embryos begin to swell to sizes comparable to normal blastocysts (Fig 5A-B, Movie 6). Single-celled embryos then form within their cytoplasm tens of fluid-filled vacuolar compartments, which can inflate into structures comparable in size to the blastocoel (Fig 5A and C, Movie 6). When measuring the growth rates of single-celled embryos we find that they are comparable to the ones of WT embryos (4.1 ± 0.8 and 3.9 ± 0.5 µm^2^/min, mean ± SEM, 4 and 20 embryos, Student’s *t* test *p* > 10^−2^, Fig 5B), indicating that they are able to accumulate fluid as fast as embryos composed of multiple cells. Therefore, single-celled embryos are able to initiate TE differentiation and display some attributes of epithelial function. Unexpectedly, this also indicates that fluid accumulation during blastocoel formation is cell-autonomous. Together, this further indicates that the developmental program entailing morphogenesis and lineage specification carries on independently of successful cleavages.

**Figure 5:**
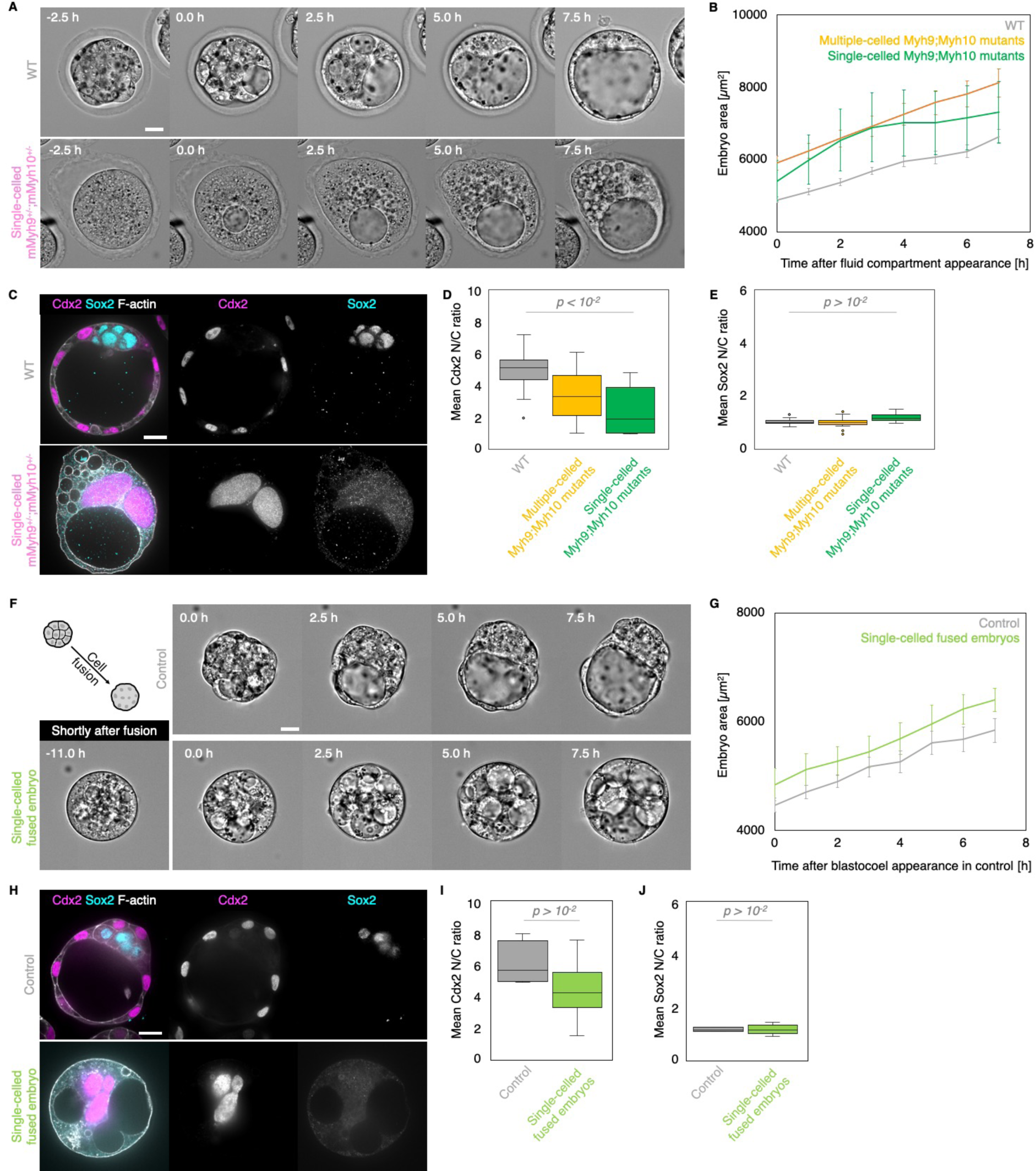
Single celled-embryos at the blastocyst stage. A) Representative images of long term time-lapse of WT and single-celled mMyh9+/−;mMyh10+/− embryos at the onset of fluid accumulation (Movie 6). B) Embryo growth curves during fluid accumulation for WT (grey, n = 20) and multiple-(yellow, n = 13)or single-celled Myh9;Myh10 (green, n = 4) mutant embryos measured for 7 continuous hours after a lumen of at least 20 µm is observed (as in blastocoel initiation of panel A). Data show mean ± SEM. C) Representative images of WT and single-celled mMyh9+/−;mMyh10+/− embryos stained for TE and ICM markers Cdx2 (magenta) and Sox2 (cyan). DAPI in yellow and F-actin in grey. The same mutant embryos as in Fig 5A are shown. D-E) N/ C ratio of Cdx2 (E) and Sox2 (F) staining for outer cells from WT (grey, n = 23) and multiple-(yellow, n = 18)or single-celled Myh9;Myh10 (green, n = 6) embryos. Mann-Whitney U test p values comparing to WT are indicated. F) Representative images of long term time-lapse of control and fused single-celled embryos at the onset of fluid accumulation (Movie 7). A schematic diagram of the cell fusion process is shown with a picture of the representative fused embryo shown right after fusion. G) Embryo growth curves during fluid accumulation for control (grey, n = 8) and fused single-celled (light green, n = 7) embryos measured for 7 continuous hours after a lumen of at least 20 µm is observed (as in blastocoel initiation of panel F). Data show mean ± SEM. H) Representative images of control and fused single-celled embryos stained for TE and ICM markers Cdx2 (magenta) and Sox2 (cyan). DAPI in yellow and F-actin in grey. I-J) N/C ratio of Cdx2 (E) and Sox2 (F) staining for outer cells from averaged control (grey, n = 4) and fused single-celled (light green, n = 6) embryos. Mann-Whitney U test p values comparing to control are indicated.

To further test whether fluid accumulation is cell-autonomous without disrupting actomyosin contractility, we fused all blastomeres at the late morula stage, before lumen formation (Fig 5F, Movie 7). The nuclei from the fused cells displayed high Cdx2 and low Sox2, which indicates that fused embryos retain a TE phenotype (Fig 5H-J). Moreover, similarly to single-celled embryos resulting from loss of contractility, single-celled embryos resulting from cell fusion grew in size at rates similar to those of control embryos while forming large fluid-filled vacuoles (3.3 ± 0.4 and 3.7 ± 0.4 µm^2^/min, 7 and 8 embryos, Student’s *t* test *p* > 10^−2^, Fig 5F-G, Movie 7). This further confirms that initiation of fluid accumulation and its rate during blastocoel morphogenesis can rely entirely on trans-cellular transport and are independent from cell-cell contacts and associated para-cellular transport.

Together, our experiments with single and double maternal-zygotic NMHC knockout embryos reveal that, while Myh9 is the major NMHC powering sufficient actomyosin contractility for blastocyst morphogenesis, Myh10 can significantly compensate the effect of Myh9 loss onto cytokinesis and compaction. Mutant and fused embryos also reveal that even blastocyst-stage embryos made out of a single cell can proceed with the preimplantation program in a timely fashion, as they differentiate into TE and build features of the blastocyst, such as the inflation of fluid-containing compartments. Finally, myosin mutants and fused embryos reveal that the developmental program entailing morphogenesis and lineage specification carries on independently of successful cleavages. This further demonstrates the remarkable regulative capacities of the early mammalian embryo.

## Discussion

Recent studies have described the critical role of actomyosin contractility in establishing the blastocyst [8,13,14,16–18]. To understand the molecular regulation of this crucial engine of preimplantation development, we have eliminated the maternal and zygotic sources of the NMHCs Myh9 and Myh10 individually and jointly. Maternal-zygotic mutants reveal that actomyosin contractility is primarily mediated by maternal Myh9, which is found most abundantly. Comparatively, loss of Myh10 has a mild impact on preimplantation development. Nevertheless, double maternal-zygotic mutants reveal that in the absence of Myh9, Myh10 plays a critical role in ensuring sufficient contractility is present to power cleavage divisions. Furthermore, double maternal-zygotic mutants bring to light the remarkable ability of the preimplantation embryo to carry on with its developmental program even when reduced to a single-celled embryo. Indeed, single-celled embryos, obtained from NMHCs mutants or from fusion of all cells of WT embryos, display evident signs of differentiation and initiate lumen formation by accumulating fluid in a timely fashion. Therefore, NMHC mutants and fused embryos reveal that the developmental program entailing morphogenesis and lineage specification carries on independently from successful cleavages.

Despite the importance of contractility during the formation of the blastocyst, maternal-zygotic mutants of NMHC had yet to be characterized. The phenotypes we report here further confirm some of the previously proposed roles of contractility during preimplantation development and provide molecular insights on which NMHC paralog powers contractility. We observe reduced compaction when Myh9 is maternally deleted and slower compaction when Myh10 is deleted (Fig 1A-B). This is in agreement with cell culture studies, in which Myh9 knockdown reduces contact size whereas Myh10 knockdown does not [30,31]. This is also in agreement with previous studies using inhibitory drugs on the preimplantation embryo. Compaction is prevented, and even reverted, when embryos are treated with Blebbistatin, a drug directly inhibiting NMHC function [13], or ML-7, an inhibitor of the myosin light chain kinase [14]. However, zygotic injection of siRNA targeting Myh9 did not affect compaction [16,41], further indicating that Myh9 acts primarily via its maternal pool. Similarly, siRNA-mediated knockdown of a myosin regulatory light chain Myl12b did not affect compaction [14]. Therefore, for efficient reduction of contractility with molecular specificity, maternal depletion is required.

The phenotypes of the maternal-zygotic NMHC mutants can appear in contradiction with conclusions from previous studies. Drugs were used to test the role of contractility on apico-basal polarity establishment with contradictory results [14]. ML-7, but not Blebbistatin, was reported to block *de novo* cell polarization [14]. Also, Blebbistatin was reported to block the maturation of the apical domain [41]. The apical domain is essential for TE differentiation and for polarized fluid transport [9,42]. We observe that even the most severely affected NMHC mutant embryos are able to establish and maintain a functional polarized fluid transport and to specifically express a TE marker (Fig 5). This argues against the requirement of contractility for de novo polarization and for apical domain maturation.

Besides compaction defects, reduced cell number is one of the major effects caused by reduced contractility in maternal-zygotic knockout of NMHCs. Loss of Myh9 alone, but not of Myh10 alone, causes cytokinesis defects (Fig 1-2, Movie 1 and 3). This could simply be explained by the higher levels of maternal Myh9 (Fig S1). In vitro studies also noted that Myh9, but not Myh10, is key to power the ingression of the cleavage furrow [27]. This recent study also noted that Myh10 can compensate the absence of Myh9 with increased cortical recruitment [27]. Consistently, we observe much more severe division failures when both Myh9 and Myh10 are maternally removed (Fig 3-5). This reveals significant compensation between NMHCs during preimplantation development. Whether this compensation takes place by a change in sub-cellular localization or by changes in expression level will require further studies.

Reduced cell number could have a major impact during very early development, when the zygote needs to produce enough cells to establish developmental patterning and morphogenesis. During normal development, the mouse embryo paces through successive morphogenetic steps of compaction, internalization and lumen formation at the rhythm of the 3^rd^, 4^th^ and 5^th^ waves of cleavages [3]. Such concomitance may initially suggest that the preimplantation program is tightly linked to cleavages and cell number. However, seminal studies have established that the mammalian embryo is regulative and can build the blastocyst correctly with either supernumerary or fewer cells [1–5]. For example, aggregating multiple embryos results in compaction and lumen formation after the 3^rd^ and 5^th^ cleavage respectively rather than when the chimeric embryo are composed of 8 and 32 cells [43]. Disaggregation leads to the same result with halved or quartered embryos compacting and forming a lumen at the correct embryonic stages rather than when reaching a defined cell number [44,45]. Contrary to disaggregated embryos, NMHC mutants and fused embryos show reduced cell number without affecting the amount of cellular material (Fig S2). The present experiments further confirm that compaction, internalization and lumen formation take place without the expected cell number. Therefore, although cleavages and morphogenetic events are concomitant, they are not linked. Furthermore, disaggregation experiments were key to reveal some of the cell-autonomous aspects of blastocyst formation: a single 8-cell stage blastomere can polarize [46] and increase its contractility [13]. Our experiments with single-celled embryos reveal that fluid accumulation as well is a cell-autonomous process (Fig 5). In fact, single-celled embryos inflate at rates similar to control embryos, implying that cell-cell contacts, closed by tight junctions at the embryo surface, are not required for accumulating fluid within blastomeres. Therefore, blastocoel components may be transported exclusively transcellularly whereas para-cellular transport through tight junctions may be negligible.

Whether single-celled embryos resulted from systematically failed cytokinesis or from cell fusion, these embryos grow at a similar rate, which is also the same as control embryos and other embryos composed of a reduced cell number (Fig 1C, 3C, 5B, 5G, S3B, S5C and S6B). This suggests that fluid accumulation is a robust machinery during preimplantation development. Interfering with aquaporins, ion channels and transporters affects fluid accumulation but the trans-cellular and para-cellular contributions remained unclear [47–50]. Future studies will be needed to understand what sets robust trans-cellular fluid accumulation during preimplantation development. Notably, fluid accumulation in single-celled embryos is accompanied by the formation of inflating vacuoles. Vacuoles were also reported to appear occasionally in mutant mouse embryos with defective cell-cell adhesion [35,51]. Similarly, vacuole formation in case of acute loss of cell-cell adhesion has been studied *in vitro* [52]. When apical lumens are disrupted, the apical compartment is internalized and this can result in fluid accumulation within vacuolar apical compartment (VAC) [52]. Analogous VACs were proposed to coalesce into the lumen of endothelial tubes upon cell-cell contact *in vivo* [53]. The molecular characteristics and dynamics of these structures and their empty counterparts, the so-called apical membrane initiation sites, have been extensively studied *in vitro* and proved helpful to better understand apical lumen formation [54,55]. In the case of the blastocyst, which forms a lumen on its basolateral side [17,50], such vacuoles may constitute an equivalent vacuolar basolateral compartment. Similarly to apical lumen, further characterization of the molecular identity and dynamic behaviors of these vacuoles may prove useful to better understand fluid accumulation during basolateral lumen formation. Studying trans-cellular fluid transport and the molecular machinery of basolateral lumen formation will be greatly facilitated by the fusion approach reported in the present study (Fig 5).

## Acknowledgements

We thank the imaging platform of the Genetics and Developmental Biology unit at the Institut Curie (PICT-IBiSA@BDD) for their outstanding support; the animal facility of the Institut Curie for their invaluable help. We thank Aurélie Teissendier for help with bioinformatic analysis, Victoire Cachoux for help with image analysis and, Julie Firmin and Yohanns Bellaïche for discussion and advice on the manuscript. Research in the lab of J.-L.M. is supported by the Institut Curie, the Centre National de la Recherche Scientifique (CNRS), the Institut National de la Santé Et de la Recherche Médicale (INSERM), and is funded by grants from the ATIP-Avenir program, the Fondation Schlumberger pour l’Éducation et la Recherche via the Fondation pour la Recherche Médicale, the European Research Council Starting Grant ERC-2017-StG 757557, the European Molecular Biology Organization Young Investigator program (EMBO YIP), the INSERM transversal program Human Development Cell Atlas (HuDeCA), Paris Sciences Lettres (PSL) “nouvelle équipe” and QLife (17-CONV-0005) grants and Labex DEEP (ANR-11-LABX-0044) which are part of the IDEX PSL (ANR-10-IDEX-0001-02). M.F.S. is funded by a Convention Industrielle de Formation pour la Recherche (No 2019/0253) between the Agence Nationale de la Recherche and Carl Zeiss SAS. Ö. Ö. is funded from the European Union’s Horizon 2020 research and innovation program under the Marie Sklodowska-Curie grant agreement No 666003 and benefits from the EMBO YIP bridging fund.

## Author contributions

M.F.S. and A.-F.T. performed experiments. M.F.S., A.-F.T., Ö. Ö., O.P., D.P., and J.-L.M. analysed the data. A.-F.T. and L.d.P. generated and maintained the mouse lines. J.-L.M. designed the project and acquired funding. J.-L.M. wrote the manuscript with inputs from all authors.

## Supplementary Figures

**Figure S1:**
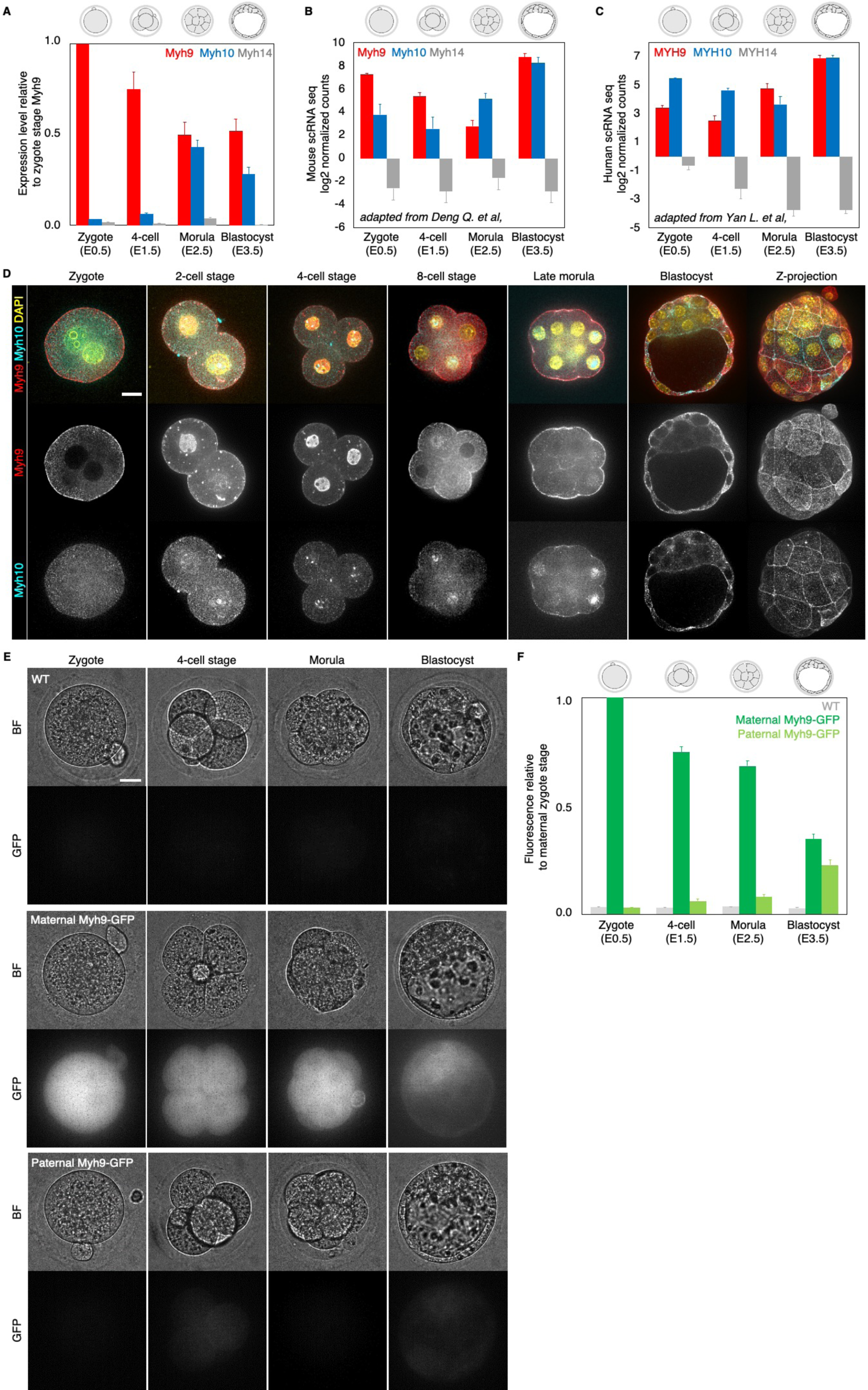
Expression of NMHC paralogs during preimplantation development. A) RT-qPCR of Myh9 (red), Myh10 (blue) and Myh14 (grey) at the zygote (E0.5, n = 234), 4-cell (E1.5, n = 159), morula (E2.5, n = 189) and blastocyst (E3.5, n = 152) stages from 6 independent experiments. Gene expression is normalized to GAPDH and shown as the mean ± SEM fold change relative to Myh9 at the zygote stage. B) Mouse single cell RNA sequencing analysis adapted from [37] showing the expression levels of Myh9 (red), Myh10 (blue) and Myh14 (grey) at the zygote, 4-cell, morula and blastocyst stages. Data show mean ± SEM. C) Human single cell RNA sequencing analysis adapted from [38] showing the expression levels of MYH9 (red), MYH10 (blue) and MYH14 (grey) at the zygote, 4-cell, morula and blastocyst stages. Data show mean ± SEM. D) Representative images of immunostaining of NMHC paralogs Myh9 (red) and Myh10 (cyan) throughout preimplantation development. DAPI in yellow. E) Representative images of WT (top), or transgenic embryos expressing a GFP-tagged version of Myh9 from their maternal (middle) or paternal (bottom) allele at the zygote, 4-cell, morula and blastocyst stages. LUT is adjusted to maternal Myh9-GFP at the zygote stage. F) Myh9-GFP fluorescent level normalized to maternal Myh9-GFP at the zygote stage. Data originate from 3 independent experiments for the WT from zygote to blastocyst stage (6 embryos), maternal Myh9-GFP zygote stage (23 embryos), maternal Myh9-GFP from 4-cell stage to blastocyst stage (24 embryos) and paternal Myh9-GFP from zygote to blastocyst stage (22 embryos). Data show mean ± SEM. Scale bars, 20 µm.

**Figure S2:**
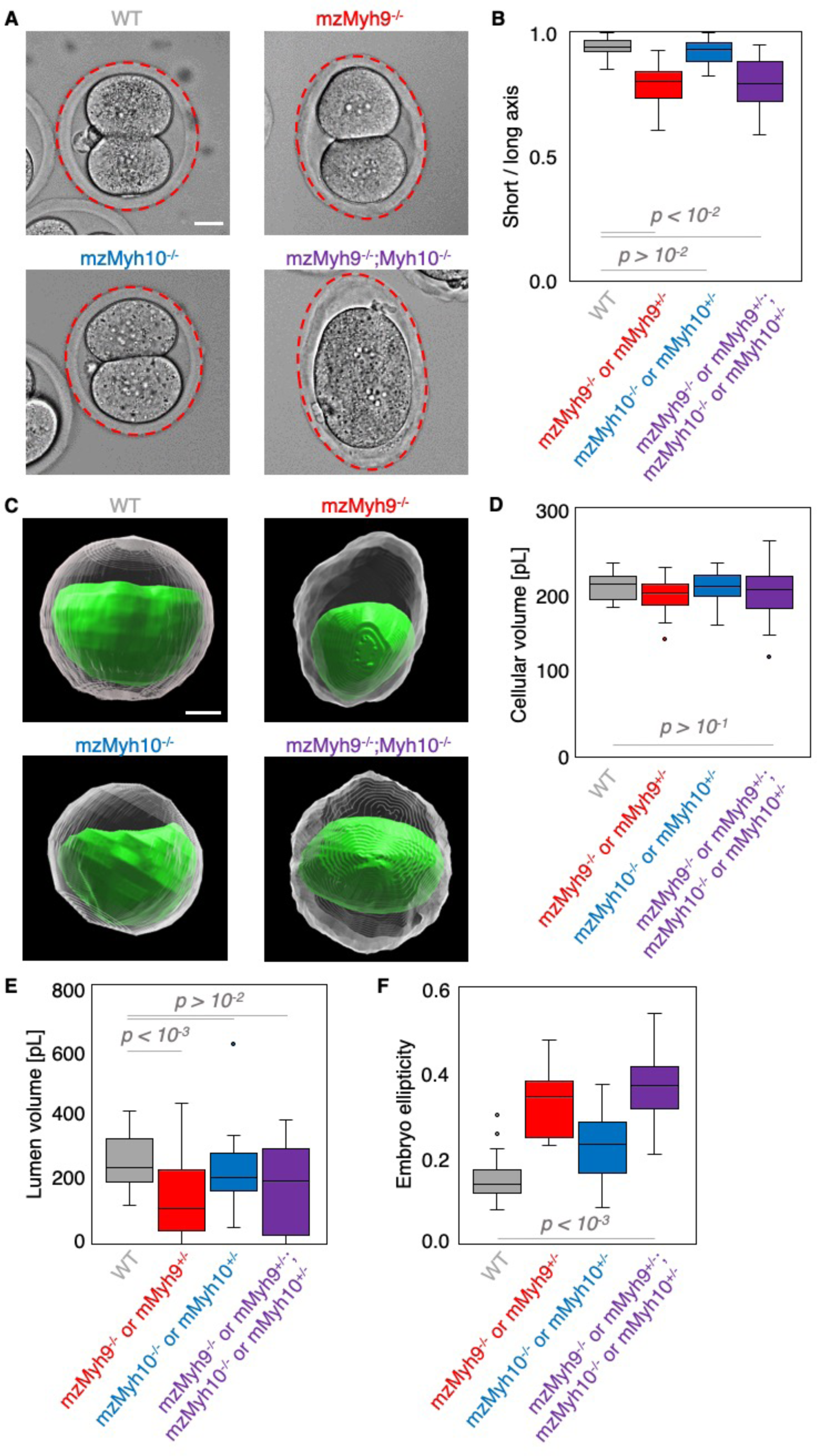
Shape analysis of maternal-zygotic Myh9 and Myh10 mutants. A) Representative images of E1.5 WT, mzMyh9−/−, mzMyh10−/− and mzMyh9−/−;mzMyh10−/− embryos. The zona pellucida is highlighted by a red ellipse. B) Boxplot of the short/long axis ratio of an ellipse manually fitted onto the zona pellucida of E1.5 embryos that are WT (n = 23), single maternal Myh9 mutants (n = 23), single maternal Myh10 mutants (n = 31) and double maternal Myh9 and Myh10 mutants (n = 25). C) Representative images of segmented WT, mzMyh9−/−, mzMyh10−/− and mzMyh9−/−;mzMyh10−/− blastocysts. The segmented lumen is shown in green. The cellular volume, shown in grey, results from subtracting the lumen volume from the segmented embryo volume. D-F) Cellular (D) and lumen (E) volumes, and 3D oblate ellipticity of segmented WT (n = 23), single maternal heterozygous and homozygous Myh9 (n = 21), single maternal heterozygous and homozygous Myh10 (n = 26) and double maternal heterozygous and homozygous for Myh9 and/or Myh10 (n = 20). Scale bars, 20 µm. Welch’s t test p values comparing to WT are shown.

**Figure S3:**
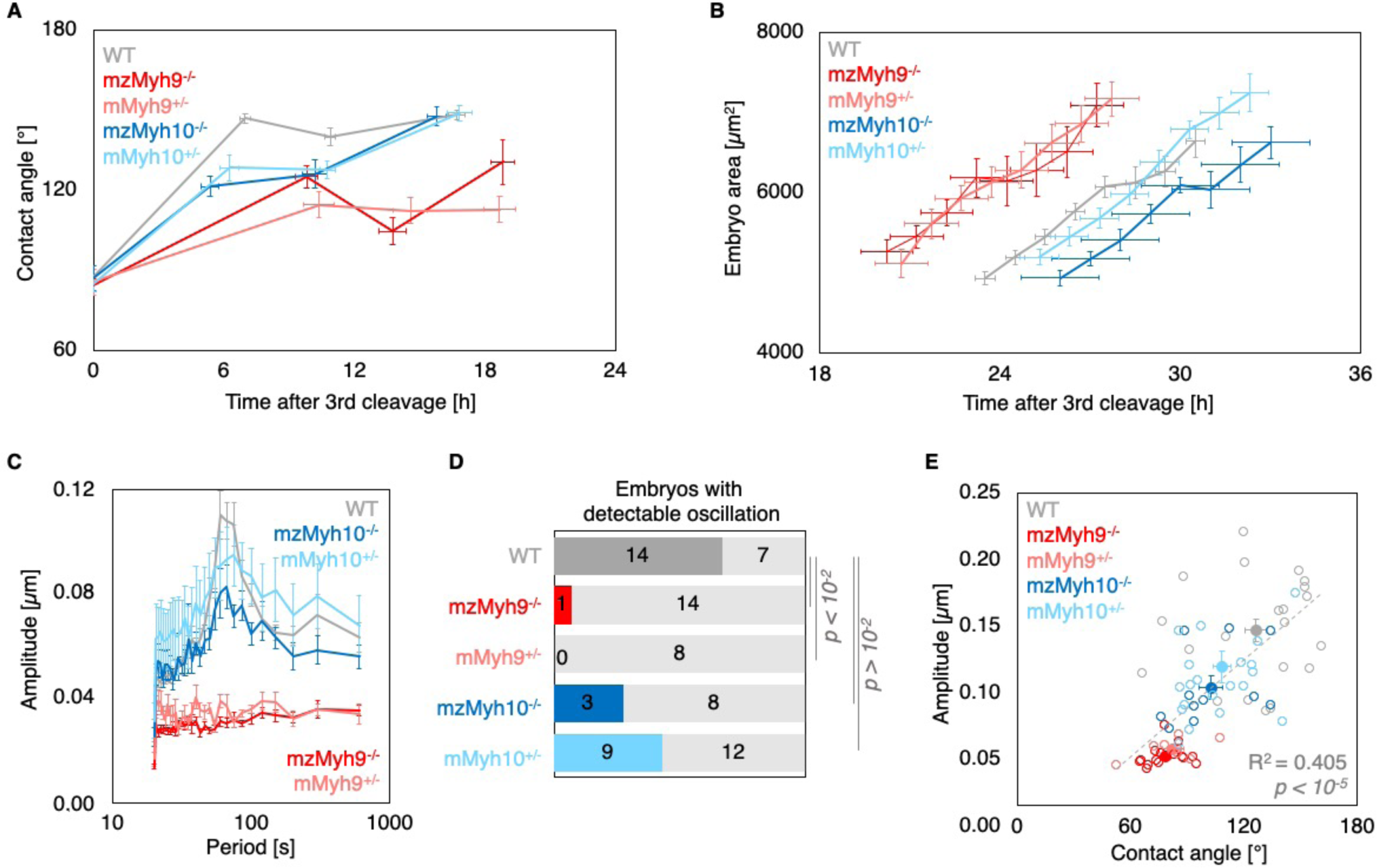
Multiscale analysis of morphogenesis in single maternal Myh9 or Myh10 mutant embryos. A) Contact angle of WT (grey, n = 23, 23, 21, 22), mzMyh9−/− (red, n = 15, 15, 10, 8), mMyh9+/− (light red, n = 8, 8, 8, 3), mzMyh10−/− (blue, n = 11, 11, 11, 11) and mMyh10+/− (light blue, n = 20, 20, 20, 20) embryos after the 3^rd^ cleavage, before and after the 4^th^ cleavage and before the 5^th^ cleavage. Data show mean ± SEM. Statistical analyses are provided in Table 1-2. B) Embryo growth during lumen formation for WT (grey, n = 20), mzMyh9−/− (red, n = 9), mMyh9+/− (light red, n = 7), mzMyh10−/− (blue, n = 7) and mMyh10+/− (light blue, n = 13) embryos measured for 7 continuous hours after a lumen of at least 20 µm is observed. Data show mean ± SEM. C) Power spectrum resulting from Fourier transform of PIV analysis of WT (grey, n = 21), mzMyh9−/− (red, n = 17), mMyh9+/− (light red, n = 8), mzMyh10−/− (blue, n = 11) and mMyh10+/− (light blue, n = 21) embryos. Data show mean ± SEM. D) Proportion of WT (grey, n = 21), mzMyh9−/− (red, n = 17), mMyh9+/− (light red, n = 8 mzMyh10−/− (blue, n = 11) and mMyh10+/− (light blue, n = 21) embryos showing detectable oscillations in their power spectrum (see Methods). Chi^2^ p value comparing to WT is indicated. E) Amplitude of oscillation as a function of the mean contact angle for WT (grey, n = 21), mzMyh9−/− (red, n = 17), mMyh9+/− (light red, n = 8), mzMyh10−/− (blue, n = 11) and mMyh10+/− (light blue, n = 21) embryos. Open circle show individual embryos and filled circles give mean ± SEM of a given genotype. Pearson’s R^2^ and p value are indicated. Statistical analyses are provided in Table 3.

**Figure S4:**
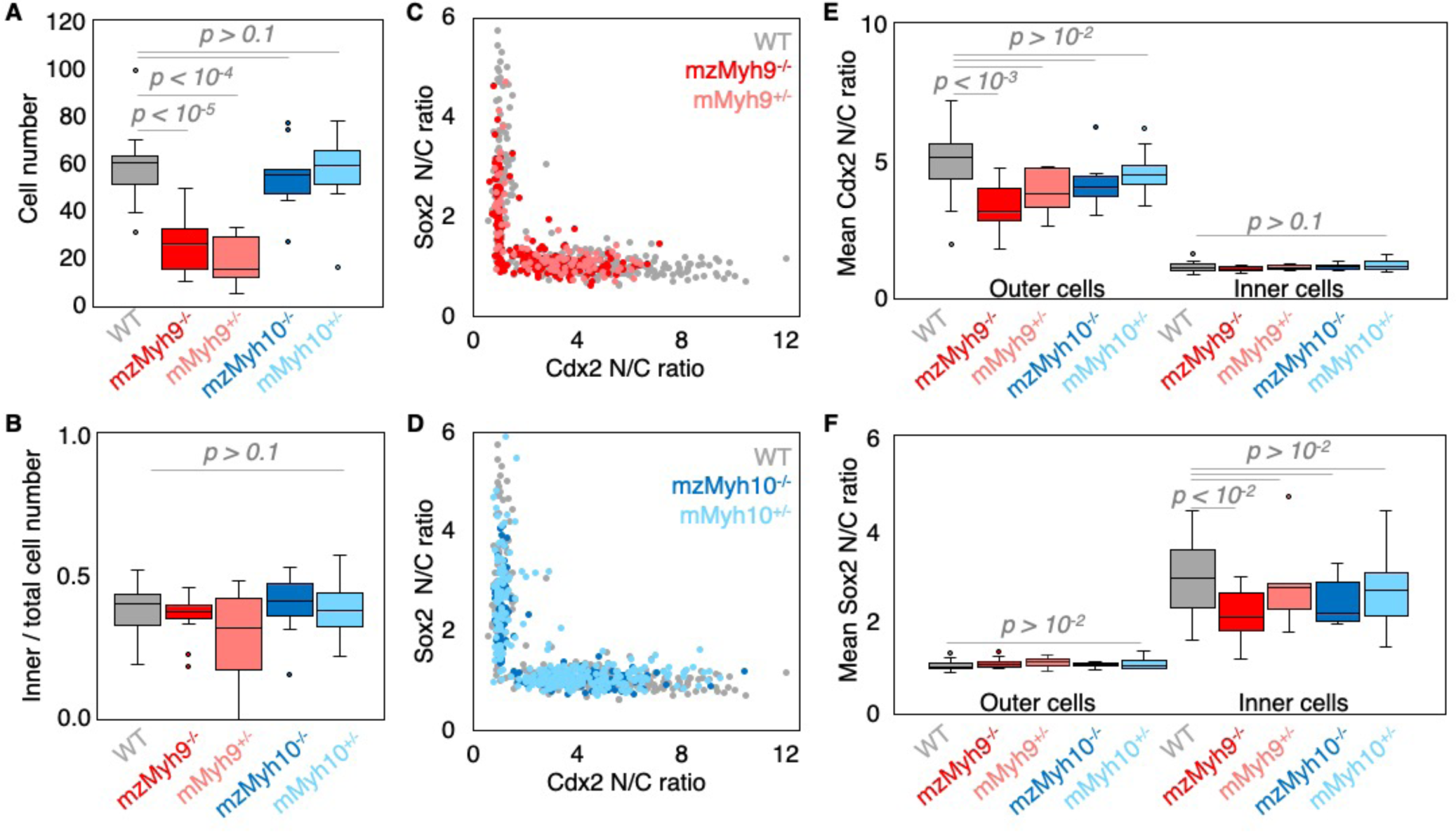
TE and ICM lineages analysis of single maternal Myh9 or Myh10 mutant embryos. A-B) Total cell number (A) and proportion of inner cells (B) in WT (grey, n = 23), mzMyh9−/− (red, n = 15), mMyh9+/− (light red, n = 8), mzMyh10−/− (blue, n = 11) and mMyh10+/− (light blue, n = 19) embryos. C-D) Nuclear to cytoplasmic (N/C) ratio of Cdx2 and Sox2 staining for individual cells from WT (grey, n = 345), mzMyh9−/− (red, n = 204), mMyh9+/− (light red, n = 95), mzMyh10−/− (blue, n = 160) and mMyh10+/− (light blue, n = 300) embryos. E-F) N/C ratio of Cdx2 (D) and Sox2 (E) staining for outer (left) or inner (right) cells from WT (grey, n = 23), mzMyh9−/− (red, n = 15), mMyh9+/− (light red, n = 8), mzMyh10−/− (blue, n = 11) and mMyh10+/− (light blue, n = 19) embryos. Mann-Whitney U test p values comparing to WT are indicated.

**Figure S5:**
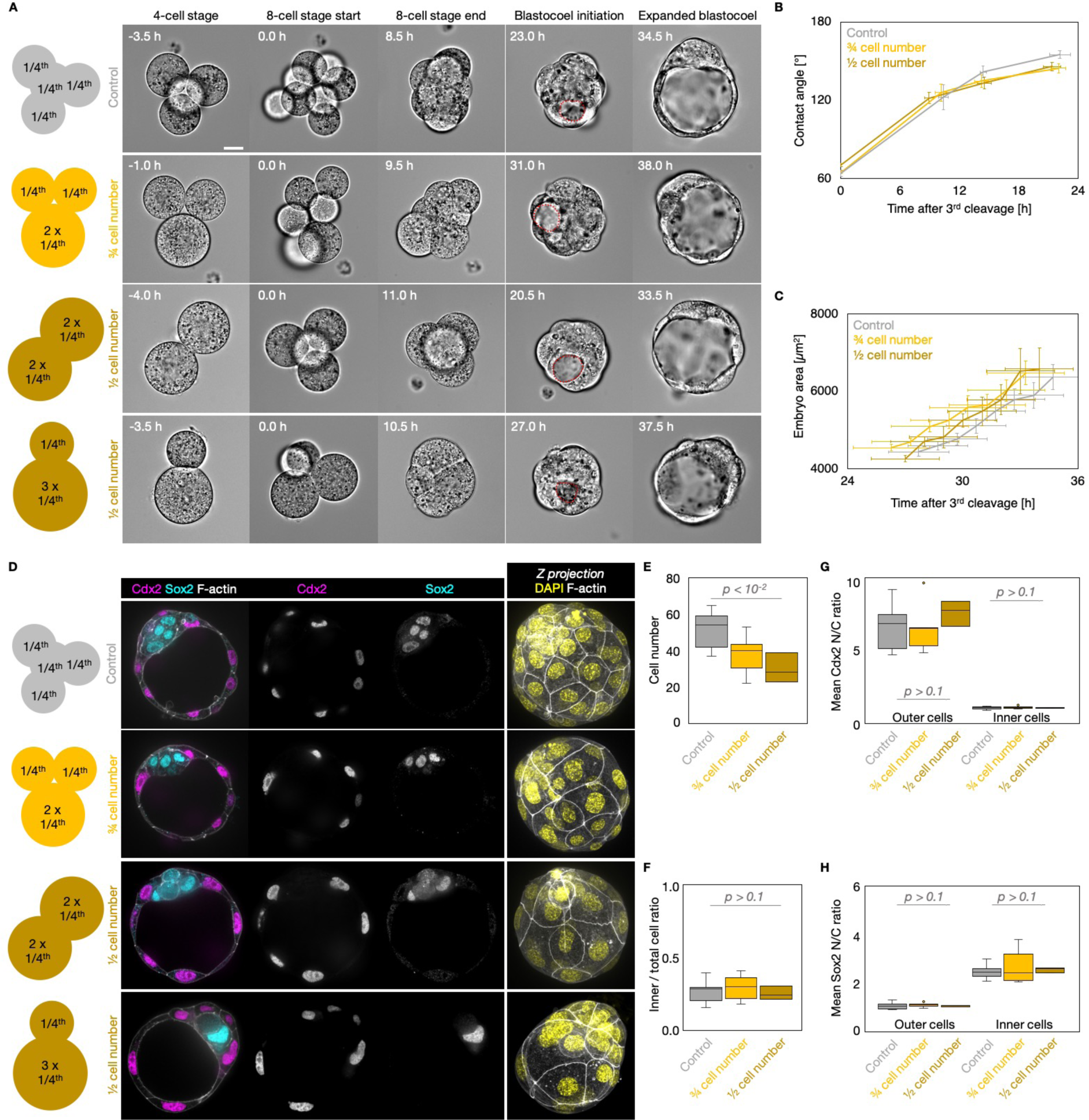
Morphogenesis and lineage specification of embryos with fused blastomeres. A) Schematic diagram of 4-cell stage embryos with 4 cells (Control) or after fusion with 3 cells, 2 cells of equal or unequal sizes. Representative images of long term time-lapse of embryos with 4 cells (Control), or after fusion with 3 cells, 2 cells of equal or unequal sizes at the end of the 4-cell stage, start and end of the 8-cell stage, at the initiation of blastocoel formation and early blastocyst stage (Movie 4). Scale bar, 20 µm. Time in hours after 3^rd^ cleavage. Dotted red lines indicate the nascent lumen. B) Contact angle of control (grey, n = 10), 3/4 cell number (yellow, n = 11) and 1/2 cell number (brown, n = 4) embryos after the 3^rd^ cleavage, before and after the 4^th^ cleavage and before the 5^th^ cleavage. Data show mean ± SEM. C) Embryo growth during lumen formation for control (grey, n = 10), 3/4 cell number (yellow, n = 11) and 1/2 cell number (brown, n = 4) embryos measured for 7 continuous hours after a lumen of at least 20 µm is observed (as in blastocoel initiation of panel A). Data show mean ± SEM. D) Representative images of embryos with 4 cells (Control), or after fusion with 3 cells, 2 cells of equal or unequal sizes stained for TE and ICM markers Cdx2 (magenta) and Sox2 (cyan). DAPI in yellow and F-actin in grey. The same fused embryos as in panel A are shown. E-F) Total cell number (E) and proportion of inner cells (F) in control (grey, n = 11), 3/4 cell number (yellow, n = 10)and 1/2 cell number (brown, n = 4) embryos. G-H) N/C ratio of Cdx2 (G) and Sox2 (H) staining for outer (left) or inner (right) cells from control (grey, n = 11), 3/4 cell number (yellow, n = 10) and 1/2 cell number (brown, n = 4) embryos. Scale bars, 20 µm. Mann-Whitney U test p values comparing to WT are indicated.

**Figure S6:**
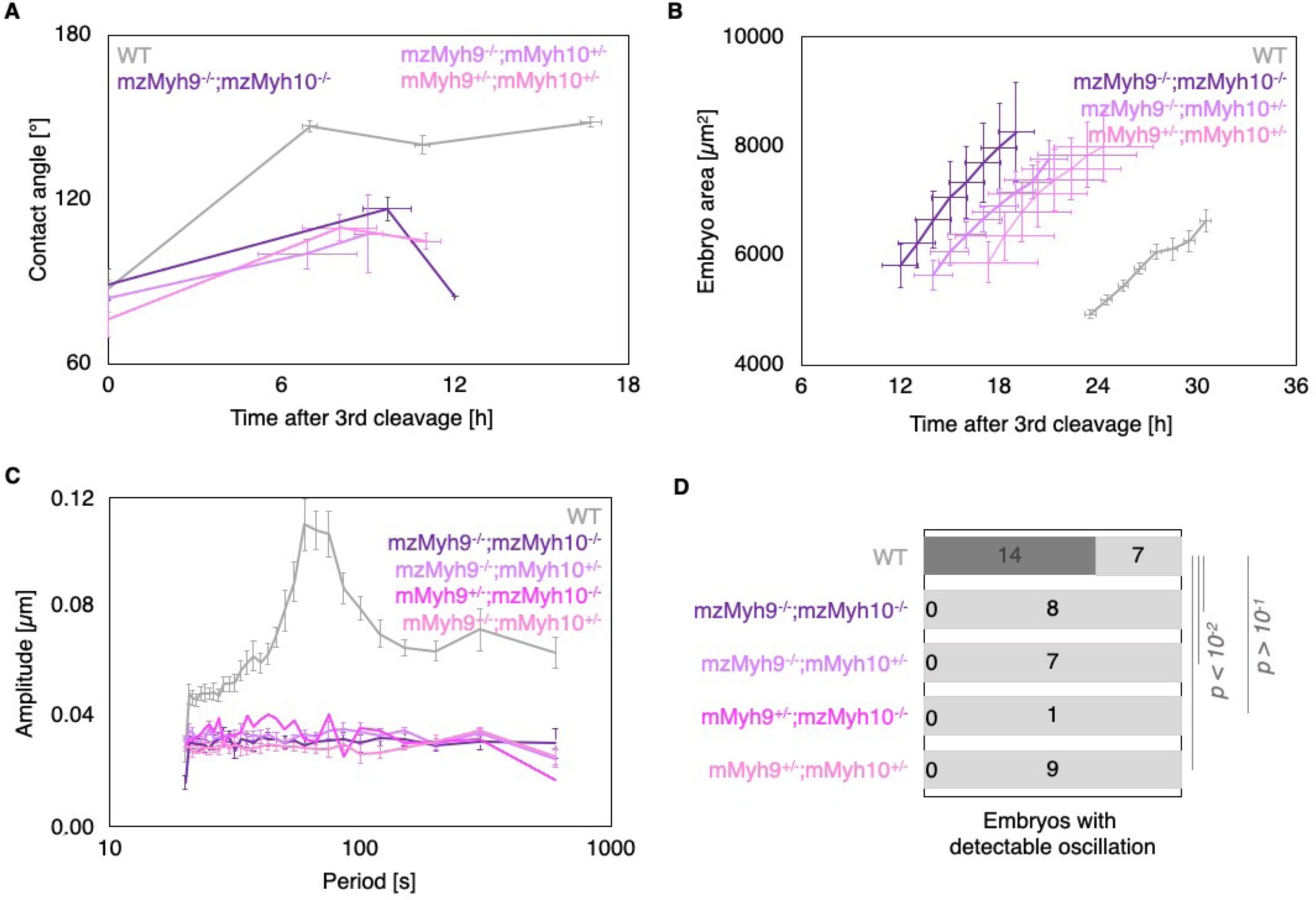
Multiscale analysis of morphogenesis in double maternal Myh9 and Myh10 mutant embryos. A) Contact angle of WT (grey, n = 23, 23, 21, 22), mzMyh9−/−;mzMyh10−/− (purple, n = 7, 3, 1), mzMyh9−/−;mMyh10+/− (lila, n = 7, 4, 2) and mMyh9+/−;mMyh10+/− (bubblegum, n = 7, 5, 3) embryos after the 3^rd^ cleavage, before and after the 4^th^ cleavage and before the 5^th^ cleavage. Data show mean ± SEM. Statistical analyses are provided in Table 1-2. B) Embryo growth during lumen formation for WT (grey, n = 20), mzMyh9−/−;mzMyh10−/− (purple, n = 5), mzMyh9−/−;mMyh10+/− (lila, n = 6) and mMyh9+/− ;mMyh10+/− (bubblegum, n = 6) embryos measured for 7 continuous hours after a lumen of at least 20 µm is observed. Data show mean ± SEM. C) Power spectrum resulting from Fourier transform of PIV analysis of WT (grey, n = 21), mzMyh9−/−;mzMyh10−/− (purple, n = 8), mzMyh9−/−;mMyh10+/− (lila, n = 7), mMyh9+/−;mzMyh10−/− (pink, n = 1) and mMyh9+/−;mMyh10+/− (bubblegum, n = 9) embryos. Data show mean ± SEM. D) Proportion of WT (grey, n = 21), mzMyh9−/−;mzMyh10−/− (purple, n = 8), mzMyh9- /-;mMyh10+/− (lila, n = 7), mMyh9+/−;mzMyh10−/− (pink, n = 1) and mMyh9+/−;mMyh10+/− (bubblegum, n = 9) embryos showing detectable oscillations in their power spectrum. Chi^2^ test p value comparing to WT is indicated.

**Figure S7:**
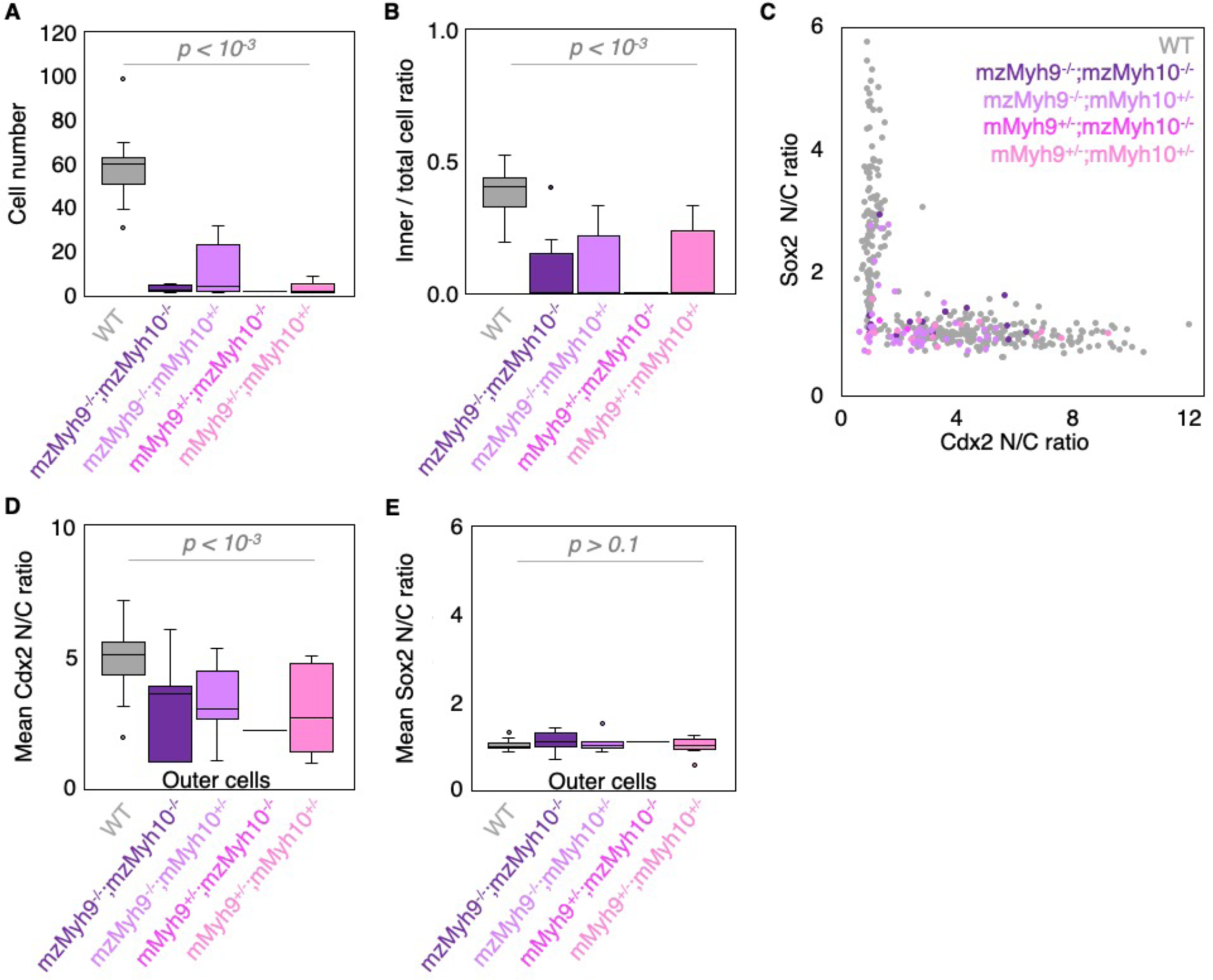
TE and ICM lineages analysis of double maternal Myh9 and My10 mutant embryos. A-B) Total cell number (A) and proportion of inner cells (B) in WT (grey, n = 23), mzMyh9−/−;mzMyh10−/− (purple, n = 8), mzMyh9−/−;mMyh10+/− (lila, n = 7), mMyh9+/−;mzMyh10−/− (pink, n = 1) and mMyh9+/− ;mMyh10+/− (bubblegum, n = 9) embryos. C) Nuclear to cytoplasmic (N/C) ratio of Cdx2 and Sox2 staining for individual cells from WT (grey, n = 345), mzMyh9−/−;mzMyh10−/− (purple, n = 17), mzMyh9- /-;mMyh10+/− (lila, n = 41), mMyh9+/−;mzMyh10−/− (pink, n = 2) and mMyh9+/−;mMyh10+/− (bubblegum, n = 21) embryos. D-E) N/C ratio of Cdx2 (D) and Sox2 (E) staining for outer cells from averaged WT (grey, n = 23), mzMyh9−/−;mzMyh10−/− (purple, n = 8), mzMyh9−/−;mMyh10+/− (lila, n = 7), mMyh9+/− ;mzMyh10−/− (pink, n = 1) and mMyh9+/−;mMyh10+/− (bubblegum, n = 9) embryos. Mann-Whitney U test p values comparing to WT are indicated.

## Movie legends

**Movie 1: Preimplantation development of WT, mzMyh9−/− and mzMyh10−/− embryos** Long term time-lapse imaging of WT (left), mzMyh9−/− (middle) and mzMyh10−/− (right) embryos from the 4-cell stage onwards. Images are taken every 30 min, scale bar is 20 µm. Time is given in hours following the last division from the 3^rd^ wave of cleavages.

**Movie 2: Periodic movements in WT, mzMyh9−/−, mzMyh10−/− and mzMyh9−/−mzMyh10−/− embryos**

Short term time-lapse imaging of WT (top left), mzMyh9−/− (top right), mzMyh10−/− (bottom left) and mzMyh9−/−;mzMyh10−/− (bottom right) embryos. Images are taken every 5 s, scale bar is 20 µm.

**Movie 3: Failed cleavage in mzMyh9−/− embryos**

Long term time-lapse imaging of a mzMyh9−/− embryo during the 3^rd^ cleavage wave. Images are taken every 30 min, scale bar is 20 µm.

**Movie 4: Preimplantation development of control, ¾ and ½ cell number embryos**

Long term time-lapse imaging of control (top left), ¾ cell number (top right) and 1/2 cell number (after fusion events of 2 pairs of blastomeres on bottom left or after fusion of 3 blastomeres on bottom right) embryos from the 4-cell stage onwards. Images are taken every 30 min, scale bar is 20 µm. Time is given in hours following the last division from the 3^rd^ wave of cleavages.

**Movie 5: Preimplantation development of mzMyh9−/−;mzMyh10−/− embryos**

Long term time-lapse imaging of mzMyh9−/−;mzMyh10−/− embryos forming an ICM (left), or without ICM (right) from the equivalent of the 2-cell stage onwards. Images are taken every 30 min, scale bar is 20 µm. Time is given in hours following the last tentative division from the 3^rd^ wave of cleavages.

**Movie 6: Preimplantation development of a mzMyh9+/−;mzMyh10+/− embryo failing all successive cleavages**

Long term time-lapse imaging of a mzMyh9+/−;mzMyh10+/− embryo failing all successive cleavages. Images are taken every 30 min, scale bar is 20 µm. Time is given in hours following the last putative division from the 3^rd^ wave of cleavages.

**Movie 7: Fluid accumulation in control and single-celled fused embryos**

Long term time-lapse imaging of control (left) and single-celled fused (right) embryos around the time of the 5^th^ wave of cleavages onwards. Images are taken every 30 min, scale bar is 20 µm. Time is given in hours following the time at which a lumen of about 20 µm is visible in the control embryo.

## Material and methods

### Embryo work

#### Recovery and culture

All animal work is performed in the animal facility at the Institut Curie, with permission by the institutional veterinarian overseeing the operation (APAFIS #11054-2017082914226001). The animal facilities are operated according to international animal welfare rules.

Embryos are isolated from superovulated female mice mated with male mice. Superovulation of female mice is induced by intraperitoneal injection of 5 international units (IU) pregnant mare’s serum gonadotropin (Ceva, Syncro-part), followed by intraperitoneal injection of 5 IU human chorionic gonadotropin (MSD Animal Health, Chorulon) 44-48 hours later. Embryos at E0.5 are recovered from plugged females by opening of the ampulla followed by brief treatment with 37°C 0.3 mg/mL hyaluronidase (Sigma, H4272-30MG) and washing in 37°C FHM. Embryos are recovered at E1.5 by flushing oviducts and at E2.5 and E3.5 by flushing oviducts and uteri from plugged females with 37°C FHM (Millipore, MR-122-D) using a modified syringe (Acufirm, 1400 LL 23).

Embryos are handled using an aspirator tube (Sigma, A5177-5EA) equipped with a glass pipette pulled from glass micropipettes (Blaubrand intraMark or Warner Instruments). Embryos are placed in KSOM (Millipore, MR-107-D) or FHM supplemented with 0.1 % BSA (Sigma, A3311) in 10 μL droplets covered in mineral oil (Acros Organics). Embryos are cultured in an incubator with a humidified atmosphere supplemented with 5% CO2 at 37°C. To remove the Zona Pellucida (ZP), embryos are incubated for 45-60 s in pronase (Sigma, P8811).

Blastomeres are fused using the GenomONE-CF FZ SeV-E cell fusion kit (CosmoBio, ISK-CF-001-EX). HVJ envelope is resuspended following manufacturer’s instructions and diluted in FHM for use. To fuse blastomeres of embryos at the 4-cell stage, embryos are incubated in 1:60 HVJ envelope / FHM for 15 min at 37°C followed by washes in KSOM. To fuse blastomeres at the morula stage, embryos are treated in the same manner in 1:50 HVJ envelope / FHM. Fusion typically completes ∼30 min after treatment. For imaging, embryos are placed in 3.5 or 5 cm glass-bottom dishes (MatTek).

#### Mouse lines

Mice are used from 5 weeks old on.

(C57BL/6xC3H) F1 hybrid strain is used for wild type (WT).

To remove LoxP sites specifically in oocytes, Zp3-cre (Tg(Zp3-cre)93Knw) mice are used [56]. To generate mzMyh9−/− embryos, Myh9^tm5RSad^ mice are used [57] to breed Myh9^tm5RSad/tm5RSad^ ; Zp3^Cre/+^ females with Myh9^+/−^ males. To generate mzMyh10−/− embryos, Myh10^tm7Rsad^ mice were used [58] to breed Myh10^tm7Rsad/tm7Rsad^ ; Zp3^Cre/+^ females with Myh10^+/−^ males. To generate mzMyh9−/−;mzMyh10−/− embryos, Myh9^tm5RSad/tm5RSad^ ; Myh10^tm7Rsad/tm7Rsad^ ; Zp3^Cre/+^ females are mated with Myh9^+/−^ ; Myh10^+/−^ males. To generate embryos with a maternal Myh9-GFP allele, Myh9^tm8.1RSad^ (Gt(ROSA)26Sor^tm4(ACTB-tdTomato,-EGFP)Luo^) females are mated with WT males; to generate embryos with a paternal Myh9-GFP allele, WT females are mated with Myh9^tm8.1RSad^ (Gt(ROSA)26Sor^tm4(ACTB-tdTomato,-EGFP)Luo^) males [59,60]. For fusion at the morula stage (Gt(ROSA)26Sor^tm4(ACTB-tdTomato,-EGFP)Luo^) mice are used [63].

#### Immunostaining

Embryos are fixed in 2% PFA (Euromedex, 2000-C) for 10 min at 37°C, washed in PBS and permeabilized in 0.1% (Sox2 and Cdx2 primary antibodies) or 0.01% (all other primary antibodies) Triton X-100 (Euromedex, T8787) in PBS (PBT) at room temperature before being placed in blocking solution (PBT with 3% BSA) at 4°C for 2-4 h. Primary antibodies are applied in blocking solution at 4°C overnight. After washes in PBT at room temperature, embryos are incubated with secondary antibodies, DAPI and/or phalloidin in blocking solution at room temperature for 1 h. Embryos are washed in PBT and imaged immediately after.

| Primary antibody | Dilution | Provider |
| --- | --- | --- |
| Cdx2 | 1:200 | Abcam, ab157524 |
| Sox2 | 1:100 | Abcam, ab97959 |
| Phospho-Myosin IIa (Ser1943) | 1:200 | Cell Signaling, 5026 |
| MYH10 | 1:200 | Santa Cruz, sc-376942 |

| Secondary antibodies and dyes | Dilution | Provider |
| --- | --- | --- |
| Alexa Fluor Plus 488 anti-mouse | 1:200 | Invitrogen, A32723 |
| Alexa Fluor 546 anti-mouse | 1:200 | Invitrogen, A11003 |
| Alexa Fluor Plus 488 anti-rabbit | 1:200 | Invitrogen, A32731 |
| Alexa Fluor Plus 546 anti-rabbit | 1:200 | Invitrogen, A11010 |
| Alexa Fluor 633 phalloidin | 1:200 | Invitrogen, A22284 |
| 4',6-diamidino-2-phenylindole (DAPI) | 1:1000 | Invitrogen, D1306 |

#### Single embryo genotyping

DNA extraction is performed on single fixed embryos, in 10 µL of DNA extraction buffer containing 10 mM Tris-HCl (pH 8, Sigma, T2694), 50 mM KCl (Sigma, 60142), 0.01% gelatin (Sigma, G1393), 0.1 mg/mL proteinase K (Sigma, P2308) at 55°C for 90 min followed by deactivation of the proteinase K at 90°C for 10 min. 2 *µ*L of this DNA extract are used in PCR reactions.

To assess the Myh9 genotype, a preamplification PCR is performed using forward (fw) primer GGGACCCACTTTCCCCATAA / reverse (rev) primer GTTCAACAGCCTAGGATGCG at a final concentration of 0.4 *µ*M. The PCR program is as follows: denaturation at 94°C 4 min; 35 cycles of 94°C 1 min, 58°C 1 min, 72°C 3:30 min, 72°C 1 min; final elongation step at 72°C 7 min. Subsequently, 2 *µ*L of the PCR product are directly used as a template for two independent PCR amplifications to detect either a 592 bp amplicon for the WT allele, with fw primer GGGACACAGTTGAATCCCTT / rev primer ATGGGCAGGTTCTTATAAGG or a 534 bp amplicon for a knockout allele, with fw primer GGGACACAGTTGAATCCCTT / rev primer CATCCTGTGGAGAGTGAGAGCAC at a final concentration of 0.4 *µ*M. PCR program is as follows: denaturation at 94°C for 4 min; 35 cycles of 94°C 1 min, 58°C 2 min, 72°C 1 min; final elongation step at 72°C 7 min.

To assess the Myh10 genotype, a preamplification PCR is performed using fw primer GGCCCCCATGTTACAGATTA / rev primer TTTCCTCAACATCCACCCTCTG at a final concentration of 0.4 *µ*M. The PCR program is as follows: denaturation at 94°C for 4 min; 35 cycles of 94°C 1 min, 58°C 1 min, 72°C 2 min, 72°C 1 min; final elongation step at 72°C 7 min. Subsequently, 2 *µ*L of the PCR product are directly used as a template for a PCR amplification using fw primer 1 TAGCGAAGGTCTAGGGGAATTG / fw primer 2 GACCGCTACTATTCAGGACTTATC / rev primer CAGAGAAACGATGGGAAAGAAAGC at a final concentration of 0.4 *µ*M. PCR program is as follows: denaturation at 94°C for 4 min; 35 cycles of 94°C 1 min, 58°C 1:30 min, 72°C 1 min; final elongation step at 72°C 7 min, resulting in a 230 bp amplicon for a WT allele and a 630 bp amplicon for a knockout allele.

### Quantitative RT-PCR

To extract total RNA, embryos are collected in 3 *µ*L of PBS, frozen on ice or snap-frozen on dry ice and stored at −80°C until further use. Total RNA extraction is performed using the PicoPure RNA Isolation Kit (ThermoFisher Scientific, KIT0204) according to manufacturer’s instructions. DNase treatment is performed during the extraction, using RNase-Free DNase Set (QIAGEN, 79254). cDNA is synthesized with random primers (ThermoFisher Scientific, 48190011), using the SuperScript III Reverse Transcriptase kit (ThermoFisher Scientific, 18080044) on all the extracted RNA, according to manufacturer’s instructions. The final product can be used immediately or stored at −20°C until further use.

All the RT-qPCR reactions are performed using the ViiA 7 Real-Time PCR machine (Applied BioSystems) according to the instruction of the manufacturer. For each target sample, amplifications are run in triplicate in 10 *µ*L reaction volume containing 5 *µ*L of 2X *Power* SYBR Green PCR Master Mix (Applied Biosystems, 4367659), 1.4 *µ*L of cDNA, 0.5 *µ*L of each primer at 2 μM and 2.6 *µ*L of nuclease free water. The PCR program is as follows: denaturation at 95°C 10 min; 40 cycles of 95°C 15 sec, 60°C 1 min; 1 cycle of 95°C 15 sec, 60°C 1 min, 95°C 15 s. The last cycle provides the post PCR run melt curve, for assessment of the specifics of the amplification. Each couple of primers is designed in order to anneal on consecutive exons of the cDNA sequence, far from the exon-exon junction regions, except for the GAPDH gene. The size of amplicons varies between 84bp and 177bp. GAPDH housekeeping gene is used as internal control to normalize the variability in expression levels of each target gene, according to the 2^-ΔCT^ method. For every experiment, data are further normalized to Myh9 levels at the zygote stage.

Total RNA was extracted from 234, 159, 189, 152 embryos at zygote, 4 cell, morula and blastocyst stage respectively, from 6 independent experiments.

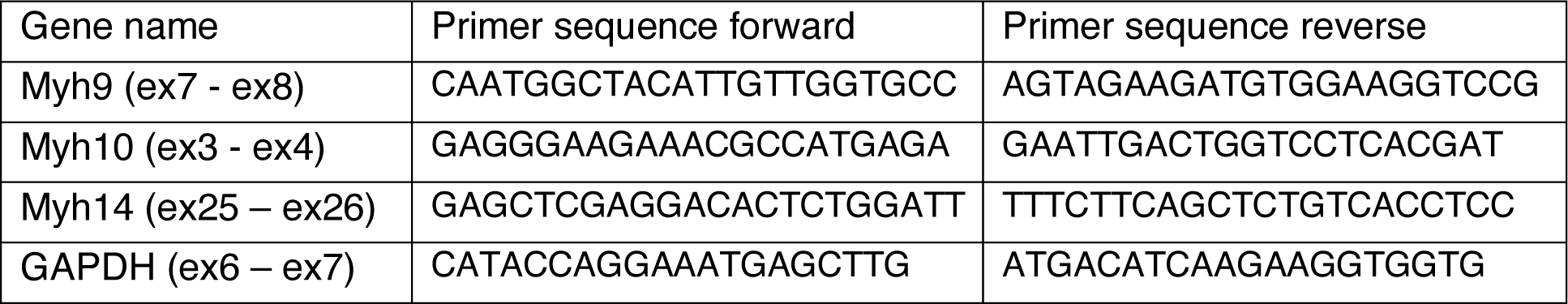

### Bioinformatic analysis

Mouse and human single cell RNA sequencing data were extracted from [37] and [38] analyzed as in [61]. In brief, single cell RNAseq datasets of human and mouse embryos (GSE36552, GSE45719) were downloaded and analyzed using the online platform Galaxy (<u>usegalaxy.org</u>). The reads were aligned to the according reference genome using TopHat (Galaxy Version 2.1.1 [62]) and read counts generated with htseq-count (Galaxy Version 0.9.1 [63]). Normalized counts were determined with limma-voom (Galaxy Version 3.38.3+galaxy3 [64]).

### Microscopy

For live imaging, embryos are placed in 5 cm glass-bottom dishes (MatTek) under a Celldiscoverer 7 (Zeiss) equipped with a 20x/0.95 objective and a ORCA-Flash 4.0 camera (C11440, Hamamatsu) or a 506 axiovert (Zeiss) camera.

Embryos are imaged from E1.5 to E3.5 until the establishment of a stable blastocoel in WT control embryos. Using the experiment designer tool of ZEN (Zeiss), we set up a nested time-lapse in which all embryos are imaged every 30 min at 3 focal planes positioned 10 *µ*m apart for 40-54 h and each embryo is subject to two 10 min long acquisitions with an image taken every 5 s at 2 focal planes positioned 10 *µ*m apart that are set 12 and 19 h after the second cleavage division of a reference WT embryo. Embryos are kept in a humified atmosphere supplied with 5% CO2 at 37°C. After imaging, mutant embryos were tracked individually for immunostaining and genotyping.

Live Myh9-GFP embryos were imaged at the zygote, 4 cell, morula and blastocyst stages. Live imaging is performed using an inverted Zeiss Observer Z1 microscope with a CSU-X1 spinning disc unit (Yokogawa). Excitation is achieved using 488 and 561 nm laser lines through a 63x/1.2 C Apo Korr water immersion objective. Emission is collected through 525/50 and 595/50 band pass filters onto an ORCA-Flash 4.0 camera (C11440, Hamamatsu). The microscope is equipped with an incubation chamber to keep the sample at 37°C and supply the atmosphere with 5% CO2.

Immunostainings are imaged on the same microscope using 405, 488, 561 and 642 nm laser lines through a 63x/1.2 C Apo Korr water immersion objective; emission is collected through 450/50 nm, 525/50 nm, 595/50 band pass or 610 nm low pass filters.

### Data analysis

#### Time-lapse of preimplantation development

Based on the time lapses of preimplantation development from E1.5 to E3.5, we assess the timing of the 3^rd^, 4^th^ and 5^th^ cleavage divisions as well as the dynamics of compaction and lumen growth following these definitions:

(1) Cleavages are defined as part of the same wave when they occur within 30 min of the cleavage of another cell within the same embryo. As embryos are being recovered at E1.5, time-lapse imaging starts around the time of the 2^nd^ cleavage. As cell divisions are affected by the mutations of Myh9 and/or Myh10, the cell number of recovered embryos is not necessarily equal to WT cell number. Therefore, we use the entire time-lapse to determine whether the first observed division corresponds to the 2^nd^, 3^rd^, 4^th^ or 5^th^ wave. Failed cytokinesis is counted as part of cleavage waves. (2) Timing of maximal compaction is defined as the time when embryos stop increasing their contact angles. (3) Timing of blastocoel formation is taken as the time when a fluid compartment with a diameter of at least 20 μm is visible.

The timings of all cleavage divisions and morphogenetic events are normalized to the end of the 3^rd^ cleavage division (beginning of the 8-cell stage in WT embryos).

Surface contact angles are measured using the angle tool in Fiji. Only the contact angles formed by two adjacent blastomeres with their equatorial planes in focus are considered. For each embryo, between 1 and 6 contact angles are measured after completion of the 3^rd^ cleavage, just before the onset of the 4^th^ cleavage, after completion of the 4th cleavage and just before the onset of the 4^th^ cleavage.

Lumen growth is assessed by the increase in area of the embryo. Using Fiji, an ellipse is manually fitted around the embryo every hour starting from the time of blastocoel formation (defined here as when a lumen of at least 20 *µ*m becomes visible) until the end of the time lapse. Growth rates of individual embryos are calculated over a 7 h time window following the time of blastocoel formation. Measured projected areas from different embryos are synchronized to the time of blastocoel formation and averaged. For fused single-celled embryo, the mean blastocoel formation time of the control embryos from each experiment is calculated to synchronize the fused embryos (1 h of deviation from the mean blastocoel formation time can be allowed to accommodate cell divisions and sampling time). For control embryos, only blastocyst growing by at least 35% of their projected area are considered.

The shape of the zona pellucida is measured by fitting an ellipse on the outer edge of the zona pellucida and measuring the long and short axes.

#### Time-lapse of periodic contractions

Particle image velocimetry (PIV) analysis is performed using PIVlab 2.02 running on Matlab [65,66]. Similar to [13], time-lapse movies acquired every 5 s are processed using 2 successive passes through interrogation windows of 20/10 μm resulting in ∼180 vectors per embryo. Vector velocities are then exported to Matlab for Fast Fourier Transform (FFT) analysis. The power spectrum of each embryo is then analyzed to assess the presence of a clear oscillation peak. The peak value between 50 and 120 s is taken as the amplitude and plotted against the mean contact angle, as this oscillation period range corresponds to the one where WT show oscillations [8,13]. An embryo is considered as oscillating when the amplitude peaks 1.7 above background (taken as the mean value of the power spectrum signal of a given embryo) to determine whether it has a detectable oscillation. Power spectra of embryos from the same genotype are averaged.

#### Time lapse of Myh9-GFP embryos

Myh9-GFP intensity is measured in FIJI by fitting a circle with a radius of 50 *µ*m over the sum projection of each embryo and measuring the mean gray value. Background intensity is measured in a circular area with radius 5 *µ*m and subtracted to the embryonic value. The corrected intensities are normalized to the zygote stage of maternal Myh9-GFP embryos.

#### Immunostaining

We use FIJI to measure the levels of Sox2 and Cdx2 expression by measuring the signal intensity of immunostainings. For each embryo, 15 cells (5 ICM, 5 polar TE, 5 mural TE) are measured. In case an embryo has fewer than 5 ICM cells, a corresponding number of TE cells are added. If the embryo has fewer than 15 cells, all cells are measured. For each cell, the signal intensity is measured in a representative 3.7 *µ*m^2^ circular area of the equatorial nuclear plane and in a directly adjacent cytoplasmic area. Mitotic and apoptotic cells are excluded from analysis. We then calculate the nuclear to cytoplasmic ratio.

To count cell number, we use DAPI to detect nuclei and phalloidin staining to detect cells. Cells are considered outer cells if they are in contact with the outside medium.

To measure the 3D properties of blastocysts, we manually segment the surface of the embryo and of the blastocoel using Bitplane Imaris. Volumes and ellipticity are obtained from Imaris. The cellular volume is obtained by subtracting the blastocoel volume from the total volume of the embryo.

#### Statistics

Mean, standard deviation, standard error of the mean (SEM), lower and upper quartiles, median, Pearson’s correlation coefficients, unpaired two-tailed Welch’s *t*-test, Mann-Whitney U-test, Pearson’s Chi^2^ test with Yates’s correction for continuity *p* values are calculated using Excel (Microsoft) and R (www.r-project.org). Pearson’s correlation statistical significance is determined on the corresponding table. Statistical significance is considered when *p* < 10^−2^. Boxplots show the median (line), interquartile range (box), 1.5 x interquartile range (whiskers) and remaining outliers (dots).

The sample size was not predetermined and simply results from the repetition of experiments. No sample was excluded. No randomization method was used. The investigators were not blinded during experiments.

### Code and data availability

The microscopy data, ROI and analyses are available on the following repository under a CC BY-NC-SA license: https://ressources.curie.fr/xxxx

